# StrataBionn: a neural network supervised classification method for microbial communities

**DOI:** 10.64898/2026.03.31.715659

**Authors:** Alex Symons, Ashley Huynh, Omar E. Cornejo

## Abstract

The classification of microbial communities into discrete states or “community state types” (CSTs) is fundamental to understanding host-microbiome interactions and their clinical implications. Traditional methods, such as the nearest-neighbor approaches, often struggle with the inherent noise, high dimensionality, and non-linear signatures of taxonomic profiles. We present a novel supervised framework for microbial community classification, leveraging an Artificial Neural Network (ANN) architecture implemented in a new tool we named StrataBionn. We rigorously evaluated our approach using large-scale vaginal microbiome datasets, directly benchmarking performance against VALENCIA and a Random Forest (RF) classifier. To demonstrate the versatility of our models, we further extended the framework to oral microbiome classification, assessing its stability across diverse anatomical sites. Our supervised models consistently outperformed the nearest-neighbor approach across all evaluated datasets. In the vaginal microbiome, our method achieved an 11.6% to 13.3% increase in performance across all primary metrics, including precision, recall, accuracy, and F1-score. Furthermore, we demonstrate that this performance advantage is maintained in the oral microbiome, highlighting the generalizability of our neural network and ensemble strategies to various microbial ecosystems without the need for niche-specific algorithmic adjustments. By capturing complex feature dependencies that distance-based methods overlook, our approach provides a more robust and accurate census of microbial community structures. StrataBionn’s ability to learn classification schemes for any microbiome with high accuracy and explainability, through the use of provided utilities to visualize feature-space classification boundaries and perform perturbation analysis on trained classifiers, makes it ideal for broad application in microecology research. This framework offers a scalable, high-performance alternative for microbiome researchers, facilitating more precise clinical stratification and biological insights across hosts body sites.

## Introduction

The study of microbial communities has fundamentally reshaped our understanding of human health, disease, and biological fitness [1–14]. It has progressed from checking for the presence of a specific taxa to tracing the shifts of community-level microbiome structures, defined by relative species abundance and functional capabilities, which more accurately describes phenotypic changes and disease risk in hosts [3,15–21]. Due to this paradigm shift, the rate at which microbiome data is generated has outpaced the ability of analysis tools to derive meaning from it. This has intensified the need for accurate, reproducible classification methods that can categorize microbiome communities to identify new biomarkers across diverse populations.

The rapid maturation of high-throughput sequencing has shown that microbial communities exhibit recurring patterns in composition across individuals, offering unprecedented opportunities to map these microbial landscapes [22–24]. However, this “big data” era presents significant analytical challenges. Extracting meaningful biological insights requires classification frameworks that are computationally efficient and capable of high-level composition analysis required in microbial ecology. Early research established the utility of composition-based community level classifications, leading to the development of standardized classification systems. Notable examples of such classifications include gut “enterotypes” [25,26] and vaginal “community state types” (CSTs) [27–29], which are consistent assemblages of microbial species, foundational in guiding modern comparative microbiome research and clinical diagnostics. Despite their utility, the methodologies used to define these categories remain a bottleneck. Historically, hierarchical clustering (HC) was the gold standard, yet its limitations in scalability and reproducibility for rapidly growing datasets are well-documented [30–36]. To address this, researchers have turned to supervised approaches like the nearest centroid classifier (NCC), implemented in robust and easy-to-use tools such as VALENCIA, which are deterministic and scale well [37]. However, NCC methods assume well-separated data and equal covariances; consequently, they often struggle with the “fuzzy”, non-linear edges and overlapping distributions typical of complex biological datasets, such as those found in microbiome studies [38,39]. Furthermore, NCC models fail to account for interactions between predictor variables—a critical oversight in microbiome data, where inter-species interactions define the community structure [40]. Finally, NCC performance decays significantly when class variances are unequal or when new data points fall outside the centroid-defined convex hulls [41,42].

Here, we present **StrataBionn**, a novel neural network-based classification algorithm designed to handle the non-linearities and high dimensionality of microbiome data. Unlike static classifiers, StrataBionn uses a neural network architecture that can be trained, saved, and adapted to a wide variety of microbial datasets. By integrating automated training methods and data preprocessing, the model achieves superior generalization and consistency. While StrataBionn offers a more sophisticated parameterization than current NCC tools, we provide comprehensive guidelines to streamline the fine-tuning process for diverse research applications.

We benchmarked StrataBionn on microbial communities from two distinct human niches: the vaginal and oral microbiomes. The vaginal microbiome is characterized by well-defined CSTs, where four types (CST-I, II, III, and V) are dominated by specific *Lactobacillus* species, and one (CST-IV) is defined by a diverse, anaerobic composition [43]. Using this established framework, we show that StrataBionn achieves higher precision and recall than existing methods. Furthermore, we applied our method to the oral microbiome, where the challenge lies in distinguishing the “oral core” from the subtle deviations associated with periodontal and systemic diseases. Finally, we show its usefulness in the assignment of samples from a novel oral sample set generated for this study.

In summary, our results indicate that StrataBionn is a highly adaptable tool that is capable of high-performance classification even with smaller reference datasets. While showcased here using vaginal and oral communities, this method is intended for broad application across any microbiome study, including comparisons between healthy and diseased cohorts. We anticipate that StrataBionn will facilitate the characterization of large-scale metagenomic datasets, ultimately deepening our understanding of the microbial drivers of host health.

## Methods

### Data Sources and Collection

#### Publicly Sourced Datasets

To train and evaluate StrataBionn, we utilized several previously published microbiota composition datasets. Vaginal microbiome profiles from **France et al.** [37] were used to train and validate the model against established Community State Type (CST) classifications, which we then validated against a dataset from **Hickey et al.** [44] to demonstrate performance on an independent dataset. Oral microbiome composition datasets were employed to evaluate the framework’s utility as a generalized classifier. As the oral microbiome research community has not established a common reference for the community types found in the oral cavity, we generated a de novo naive classification with a large dataset recently generated by **Manghi et al.** [45]. We then used the naive classification as “true” labels for the training and validation of our method. We show that this approach produces similar results, in terms of performance metrics, to those obtained in a more curated dataset like the vaginal microbiome. We show how the classifier can be used to classify samples on a sample set generated for this work and a dataset from published studies [46]. A freeze of the public datasets used for the study are available in the github repository of the tool.

#### In-House Oral Microbiome Collection and metagenomic sequencing

To test the classification consistency in the oral microbiome, plaque samples were collected from adult refugees (≥18 years) at the Rwamwanja UN refugee settlement, Uganda. Ethics and Recruitment: Collection was approved by the Institutional Review Board at Washington State University (IRB #15196-002). A total of 54 participants were recruited using a random number sequence generated in R to ensure unbiased selection. Informed consent was obtained from all participants and a translator was present when necessary. Sampling and Storage: Dental plaque was collected during routine cleanings, preserved in RNAlater, and stored at −80°C after transport to Pullman, WA, under CDC Import Permit #2016-03-212. Library Preparation and Sequencing: Microbial DNA was extracted using the MoBio Powerlizer™ DNA Isolation Kit (Mo Bio Laboratories, Carlsbad, CA, USA), following the manufacturer’s protocol. For this, plaque samples preserved in RNAlater were spun down, RNA later discarded and macerated manually with sterile pestles with lysis solution. After DNA extraction, samples were sheared using a Covaris M220 to a 550 bp insert size. Libraries were prepared with the NEBNext Ultra DNA Library Prep kit (New England Biolabs Inc.) and sequenced on a single lane of an Illumina HiSeq 2500 at the Washington State University Genomics Core.

#### Bioinformatics and Quality Control of novel oral metagenomic data

Initial quality assessment was performed using FastQC [47]. Out of 54 samples, 39 produced enough DNA after extraction and/or passed FastQC’s quality control test, and were used in analysis. Sequence trimming was performed with TrimGalore and Cutadapt and we hard trimmed the first 11 bp on the 5’ end and soft trim for Phred scores < 25 [48,49]. Host Filtering: reads were aligned to the human reference genome (GRCh38) using STAR [50], and human reads were removed and deleted permanently. Taxonomic Classification: non-human reads were classified using KrakenUniq [51], and to minimize false positives, we required ≥100 unique kmers and a duplicity ratio ≤ 3. Count matrices were generated at the genus and species levels. Raw sequence data is available at SRA-NCBI through accession PRJNA1445365.

### Microbiome classifier: the StrataBionn Framework

We developed StrataBionn, stratification of biological data using neural networks, a supervised classification tool designed to assign CST, or other community level labels, to microbiome data. It accepts as an input a standardized data format (CSV) similar to that used by VALENCIA, which includes columns for sample ID, read counts, labels, and a column for each taxa for which count data was collected. The method incorporates a stratified data partitioning strategy to ensure robust model training and evaluation, particularly in scenarios with varying data availability.

#### Stratified Data Partitioning

To maintain the inherent class balance of the original labeled data within each subset and mitigate potential biases, a stratified splitting approach was employed. This ensured that the amount of each CST in each subset used during training is consistent with the distribution found in the original, unpartitioned dataset, to avoid sampling biases during model training. Datasets were first shuffled and then partitioned proportionally by CST label. Specifically, for each CST, the designated proportions of samples were allocated to the training, testing, and validation sets. Following the split, the proportional distribution of CST labels within each subset was recalculated and compared to the original dataset distribution. Consistency was automatically assumed if the proportional representation of each CST in the resulting subsets was within a predefined tolerance of 0.1% of the original distribution. We defined tolerance using the equation 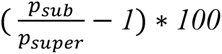, where *p_sub_* is the proportion of a given CST in the subset of interest (training, test, validation), and *p_super_* is the proportion of that same CST in the superset which contains all the training, testing, and validation samples. This rigorous stratification ensures that each data subset provides a representative snapshot of the overall class distribution, facilitating unbiased model training and evaluation. Two data partitioning schemes were implemented and evaluated: (i) a 80/10/10 split for training, testing, and validation sets, respectively, simulating conditions with ample data; and (ii) a 60/20/20 split for training, testing and validation of the same sets, mimicking data-limited scenarios. The processed and partitioned datasets were then exported as comma-separated value (CSV) files for subsequent model training. These files were formatted in the same manner as those accepted by VALENCIA, containing columns for the sample id, total number of reads, sample label (when training), and bacteria species. This format was chosen to increase compatibility between StrataBionn and other existing tools (i.e. VALENCIA).

#### Data Preprocessing within StrataBionn

Prior to model training, StrataBionn performs several crucial preprocessing steps. To account for variations in sequencing depth or sample “quality,” the raw feature counts were normalized by the total counts per sample. This transformation ensures that samples with differing sequencing depths contribute equally to the model training process. Subsequently, any null or zero values, which could lead to computational errors during training, were imputed with a small, non-interfering numerical value. While the StrataBionn framework includes the option for additional data normalization techniques, initial evaluations indicated that the total count normalization alone yielded the most effective model performance for this specific application, and thus, further normalization steps were not performed in the final model training and evaluation.

#### Classification Algorithms

Two distinct supervised learning algorithms were implemented and evaluated within the StrataBionn framework: a Random Forest Classifier (RFC) and an Artificial Neural Network (ANN). These methods were selected for their classification consistency and evaluated along with the VALENCIA classifier which served as an accuracy baseline.

- **Random Forest Classifier:** For this model, we used the RandomForestClassifier module from the python scikit-learn library [52]. The RFC was parameterized to optimize its performance. The number of estimators (n_estimators) and the number of features to consider when looking for the best split (max_features) were systematically tuned. Numbers of estimators ranging from 100 to 10,000 were tested, and while training time increased significantly with more estimators, performance plateaued around 95.1% (10,000 estimators), taking around 7 minutes.
- **Artificial Neural Network:** The ANN model consisted of an input layer, a single hidden layer, a Leaky Rectified Linear Unit (Leaky ReLU) activation function following the hidden layer, and a dropout layer for regularization. The size of the hidden layer was parameterized, with a default setting of 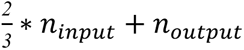, where *n_input_* represents the number of input neurons and *n_output_* the number of output neurons. The equation for the full system is as follows: 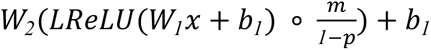, where *x* is the input data vector, *W_1_*, *W_2_* are learnable weight matrices, *b_1_*, *b_2_* are learnable bias matrices, *p* is the probability that a neuron will dropout, and *m* is a vector that masks neurons to be dropped via multiplication with 0 or 1. The optimization algorithm was selectable between Adaptive Moment Estimation (ADAM) and Stochastic Gradient Descent (SGD). The SGD optimizer offered an optional class weighting scheme to address potential imbalances in CST prevalence, allowing for increased emphasis on accurately classifying less frequent classes. The learning rate and a learning rate reduction factor were also tunable hyperparameters. A probability of assignment is reported for all CSTs for each sample, and the CST with the highest probability is assigned as the label for each sample. The model was constructed using the pytorch library and written python [53]. Training time for the vaginal microbiome test set with default hyperparameters took around 27 seconds.
- **VALENCIA Baseline:** The previously established VALENCIA method was included as a baseline for performance comparison. It uses a nearest centroid classification model, which employs pre-provided centroids constructed to classify vaginal microbiome samples.

We also provide a naive assessment of the uncertainty in the classification by extracting the logits from the softmax assignment and standardizing those values to generate probabilities of samples belonging to the assigned class. For this we exponentiated the raw logits and divide them by the sum of all exponentiated values according to the following formula, as implemented in the pytorch library [53]:

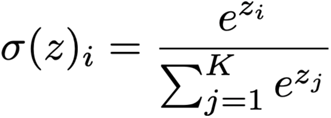

### Evaluation and Validation Strategy

#### Model Evaluation

All trained models, including the baseline VALENCIA classifier, were evaluated using the same vaginal microbiome dataset. Critically, each proposed model was evaluated on a distinct test set that was not used during training or validation, ensuring an unbiased assessment of generalization performance. The VALENCIA baseline was evaluated using its provided centroids, which were constructed based on the entire dataset. Model performance was quantified using classification accuracy, defined as the proportion of correctly classified samples.

#### Evaluation as General Microbiome Classifier

Performance of the ANN on oral microbiome datasets was measured to evaluate its usefulness as a generalized microbiome community classification method. We first performed a naive classification of the large oral microbiome dataset using K-Means clustering to generate labels that would be employed for model training and evaluation. Oral data was reformatted and treated identically to the vaginal microbiome data, using the same processing methods previously described. Evaluation of the ANN on oral microbiome data was performed using the same classification accuracy measurement used for the vaginal microbiome. The Random Forest Classifier was not evaluated due to its much longer training time to achieve a lower accuracy compared to the ANN, and VALENCIA was not evaluated due to its specificity to the vaginal microbiome. We analyzed the performance of the method with the oral microbiome.

#### Application of Trained Classifiers Across Studies

A vaginal microbiome test set containing population data on different bacteria species [44] was classified using a model trained to identify the CSTs from the VALENCIA training set. StrataBionn was first trained on the VALENCIA training set while limited to considering only the set of bacteria species common to both the training dataset and the target dataset. The trained classifier was then used to apply CST labels to all samples in the target dataset, ignoring bacteria species which are not part of the common set. These CST classifications were then manually examined and determined to be reasonable based on their bacterial makeup using the findings of France et al [37].

## Results

### Training, Validation and Test datasets

For the vaginal Microbiome, we generated stratified random sets from the France et al [37], for training (80%: 10,580; 60%: 7,935), validation (80%: 1,334; 60%: 2,655), and testing (80%: 1,317; 60%: 2,641). For the oral microbiome, we generated stratified random sets from the Manghi et al. [45] study for training (80%: 6,248; 60%: 4,686), validation (80%: 784; 60%: 1,565), and testing (80%: 780; 60%: 1,561).

We obtained read-counts supporting the presence of bacteria species in 39 newly generated oral microbiomes for this study. The number of reads and estimated relative abundances for species can be found in the github repository.

### Preprocessing of Vaginal Microbiome Datasets

During the development and testing of StrataBionn, two training data sets with 80% and 60% training allocations were included to test scenarios when training data is abundant or limited. Stratified datasets from France et al. [37] containing curated CST classifications were used for their high quality labels and large number of samples (13,231). The PACMAP [54] representations of the truth labels for the high training data availability scenario are shown in Figure 1, demonstrating similar CST composition among subsets. Consistent with our expectations, more closely related CST communities are clustered closely in the PACMAP representation (Figure 1). For example, CST-IA and CST-IB, which both feature a high abundance of *Lactobacillus crispatus*, have a similar projection in the map and tend to occur more closely than less related state types, such as CST-IA and CST-V. This observation supports our use of PACMAP projections of the data for the purpose of representation.

**Figure 1.**
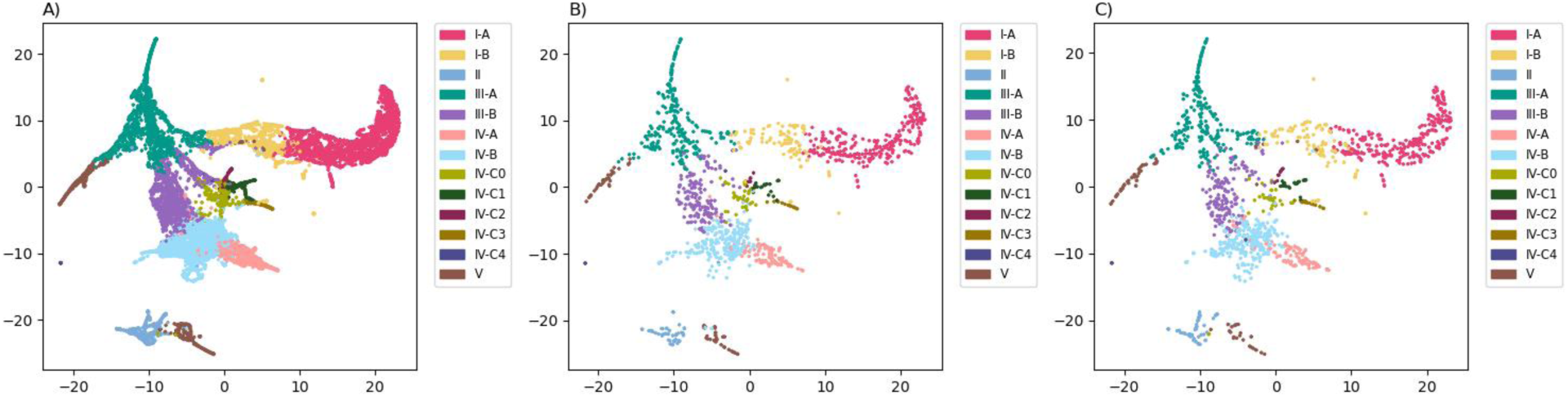
PACMAPs showing classifications from France et al. [37] for A) the 80% training set, B) the 10% testing set, and C) the 10% validation set.

The worst CST distribution tolerances across all CSTs are 0.005%, 7.2%, and 7.2% for the 80% training (Figure 1A), 10% testing (Figure 1B), and 10% validation (Figure 1C) sets respectively. For the 60% training, 20% testing, 20% validation sets, the tolerances are 0.05%, 13.4%, and 13.4% respectively. PACMAP representation of the 60%, 20%, 20% configuration is presented in the supplementary materials (Supplementary Figure 1).

### Evaluation of Classification Models

Each classification method was evaluated on both the high and low training data availability labeled datasets using accuracy (Figure 2A), recall (Figure 2B), F1 score (Figure 2C), and precision (Figure 2D) as metrics. These metrics were selected to measure the number of samples correctly identified (accuracy), the rate of false negatives (recall), the rate of false positives (precision), and balanced measure of both recall and accuracy (F1). Evaluation was performed on the validation dataset after training a classifier on the corresponding training dataset for all classifiers except for VALENCIA, where the original author provided centroids were used for classification [37]. While the ANN and RF classifiers trained on a subset of the Vaginal microbiome data, VALENCIA’s centroids were generated using the full microbiome dataset as described in the author’s methodology [37]. For the ANN classifier, probabilities of assignment were above 87% for 95% of samples (80% training: 87.8%; 60% training: 87.6%), and around 95% for 80% of classified samples (80% training: 95.3%; 60% training: 94.9%). In all cases the neural network classifier and random forest classifier outperformed VALENCIA when classifying both datasets. The ANN achieved a **11.6-13.3% increase** compared to VALENCIA across all metrics (Figure 2A-D), performing slightly better with higher data availability, while the Random Forest approach achieved a 9-10.8% increase across all metrics compared to VALENCIA (Figure 2A-D). The Random Forest classifier performed slightly better in lower data availability scenarios, but never outperformed the ANN classifier by any measured metric.

**Figure 2.**
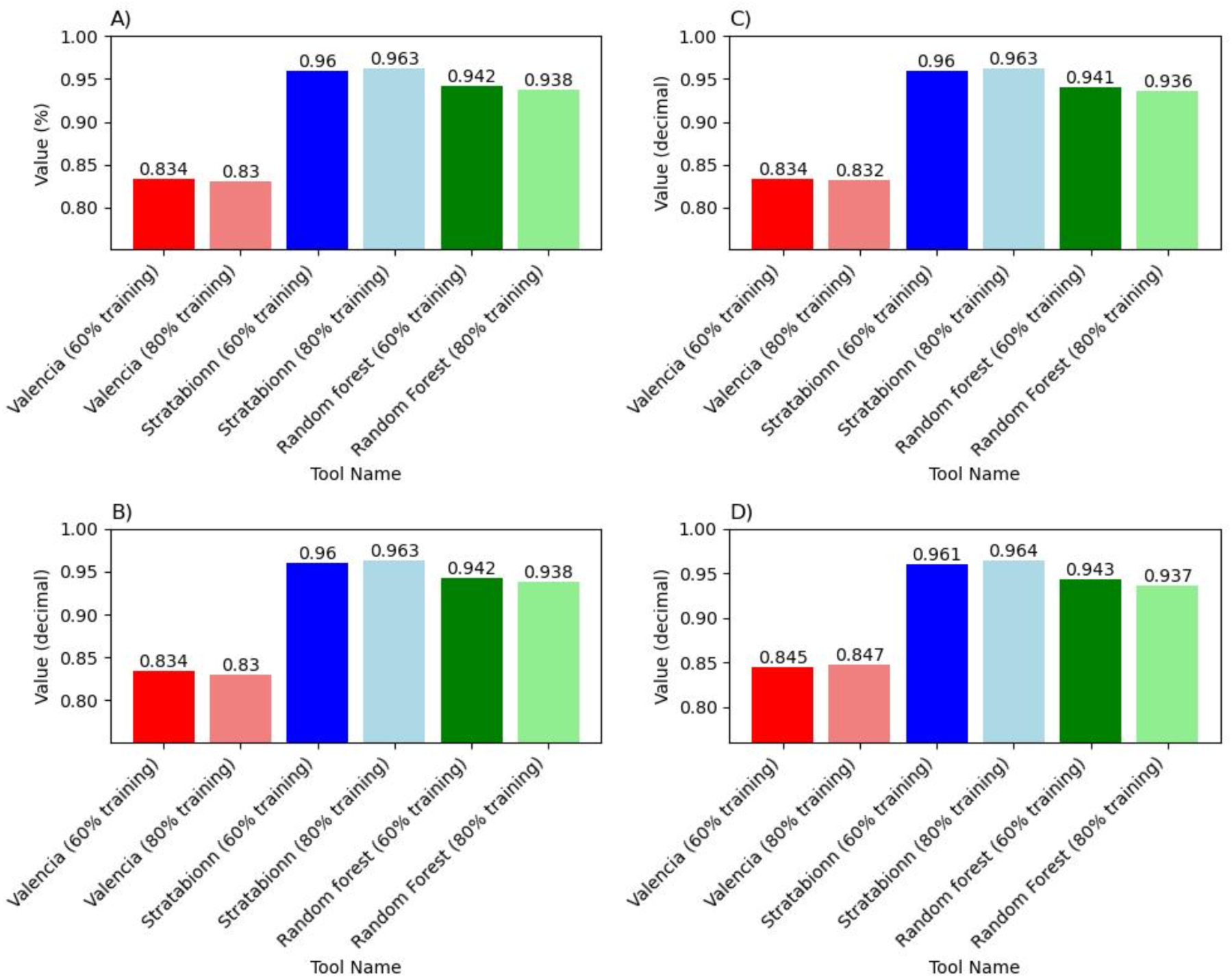
Precision metrics (A) Accuracy, (B) Recall, (C) F1-score, and (D) Precision for Valencia, StrataBionn (60% training), StrataBionn (80% training), Random Forest (60% training), Random Forest (80% training).

To understand the conditions which led to our models’ increased performance, we generated confusion matrices for each classification method by comparing the validation dataset truth values to StrataBionn-assigned CST labels. In VALENCIA, misidentification of CST-IA, CST-IB, CST-IIIA, CST-IIIB, and CST-IVA disproportionately contributes to its lower performance (Figure 3A). In the case of the ANN classifier and random forest classifier implemented in StrataBionn, communities CST-IA and CST-IIIA disproportionally contribute to the misassignment of the community labels; while CST-IIIB and CST-IVB marginally contribute to the misassignment (Figures 3B-C). To better demonstrate the cases where StrataBionn outperforms VALENCIA in assignments, we estimated the differences in the number of classified communities for each cell in the confusion matrix (Figure 3D). Cases where the predicted label matched the true label were observed to be higher for the neural network classifier compared to VALENCIA for most CSTs, especially those which are less common (blue diagonals in Figure 3D). Hotspots denoted in red on the off-diagonal of the comparative confusion matrix show that the neural network classifier is better able to differentiate between some CSTs with similar composition distributions than VALENCIA (Figure 3D). This is likely due to the ability to classify along non-linear boundaries which is not possible using a nearest centroid classifier based approach. Due to the greater performance and higher ability to differentiate between similar CSTs, we chose to proceed with a neural network classifier as the basis for StrataBionn.

**Figure 3.**
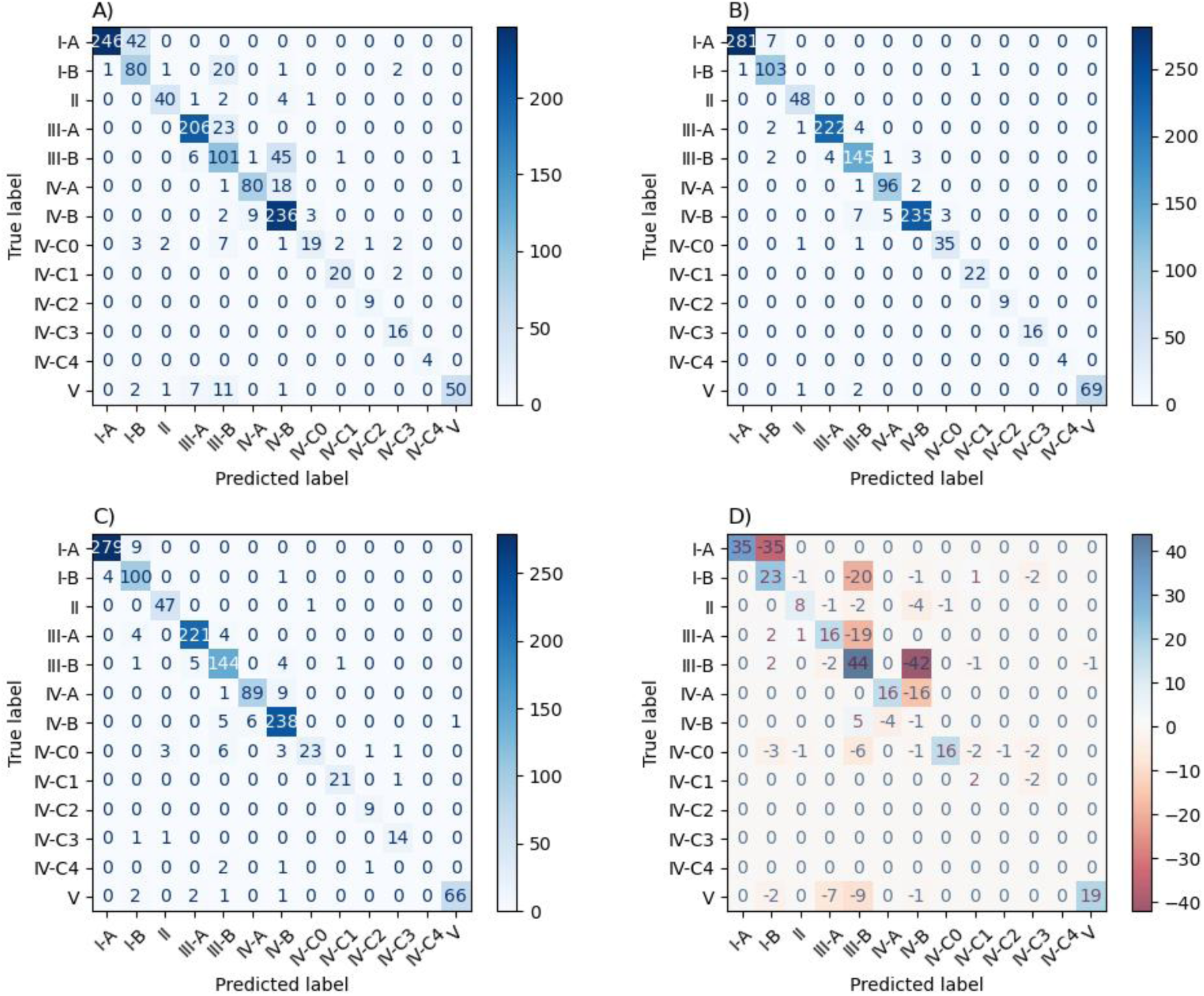
Confusion matrices for: (A) VALENCIA (classifying the 80% training set), (B) StrataBionn (80% training), (C) Random Forest (80% training), and (D) Differential between StrataBionn minus VALENCIA scores. In all figures the counts on the diagonal correspond to successful assignments of communities by the methods to their true labels and the off-diagonals correspond to misassignments. In figure 3D the positive values correspond to a larger number of assignments to that category by StrataBionn, while negative numbers correspond to a lower number of assignments by StrataBionn. Almost all elements on the diagonal of Figure D are positive and all in the off-diagonal are negative, indicating that StrataBionn outperforms VALENCIA in assigning communities to their true label, while reducing the misassignments. The only exception is one fewer true assignment to CST IV-B compared to VALENCIA.

### Evaluation of Novel Dataset Classifications

To test the ability of StrataBionn to apply accurate classifications to novel datasets, we generated classifications for an unclassified vaginal microbiome dataset containing 458 samples from Hickey et al. [44]. This dataset required no preprocessing other than reformatting and column renaming. The classification model we used for this task was trained on the labeled France et al. [37] dataset, and configured to consider only the bacteria species common to both datasets. We then applied this classifier to the full Hickey et al. [44] dataset to get CST assignments for each sample. We generated two PACMAPs using both the classified Hickey et al. [44] dataset and the truth labels from the France et al. [37] dataset, shown in figure 4. We observed that the novel classifications were assigned labels that matched their surrounding samples with similar compositions. Assigned samples cluster around data points with the same classification from the original France et al. [37] dataset.

**Figure 4.**
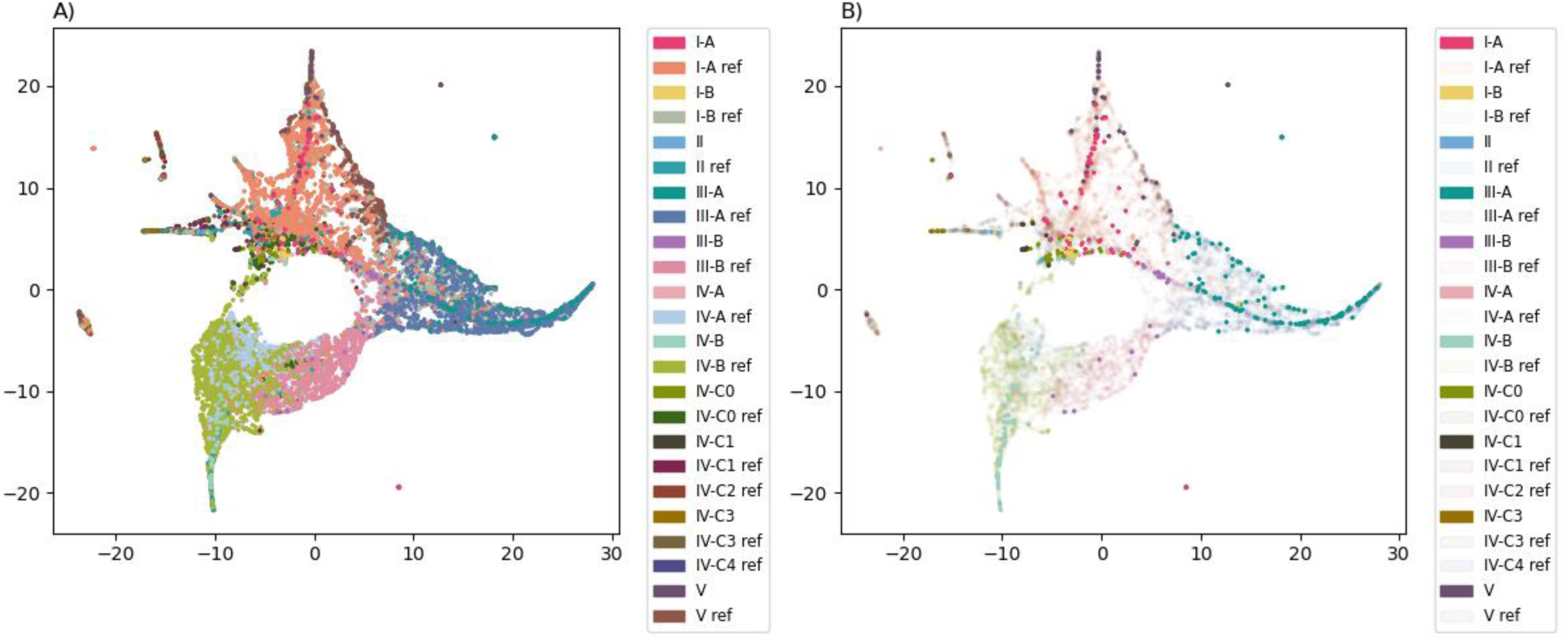
PACMAPs of reference dataset classifications and Hickey et al. [44] dataset classifications generated by StrataBionn. In figure (A), both datasets are displayed with equal opacity. In figure (B), classified Hickey et al. [44] data is displayed at full opacity while reference data is displayed at 5% opacity.

### Classification of Oral Microbiome Datasets

An oral microbiome bacterial community composition dataset was used to validate StrataBionn’s ability as a general purpose classification tool on non-vaginal microbiome datasets. We used 7,812 samples collected by Manghi et al. [45] with pre-existing classifications removed due to poor correlation with the data (see supplementary figure 2). As oral microbiomes have not been characterized to the same level of detail of classification as the vaginal microbiomes, we employed a K-Means clustering algorithm to determine the optimal number of clusters for this dataset. We found that using 3 clusters provided the optimal classification by overserving the silhouette and elbow plots provided in supplementary figures 3 and 4. After clustering labels were applied, the data was then partitioned into high and low data training data availability datasets using the stratified partitioning process previously described for the vaginal microbiome. PACMAP visualizations of these subsets are provided in Figure 5. The worst CST distribution of tolerance values for these datasets, as defined by the percent occurrences in a set above (positive) or below (negative) the occurrences found in the reference set, were 0.008%, 0.0074%, and 0.0074% for the 80% training (Figure 5A), 10% testing (Figure 5B), and 10% validation (Figure 5C) sets respectively. For the 60% training, 20% testing, 20% validation sets, the tolerances were 0.019%, 0.35%, and 0.35% respectively. We observed that samples which were assigned to the same cluster exhibit spatial locality, suggesting that our assignments capture the composition of the samples, making them ideal for evaluating the performance of our model (Figure 5).

**Figure 5.**
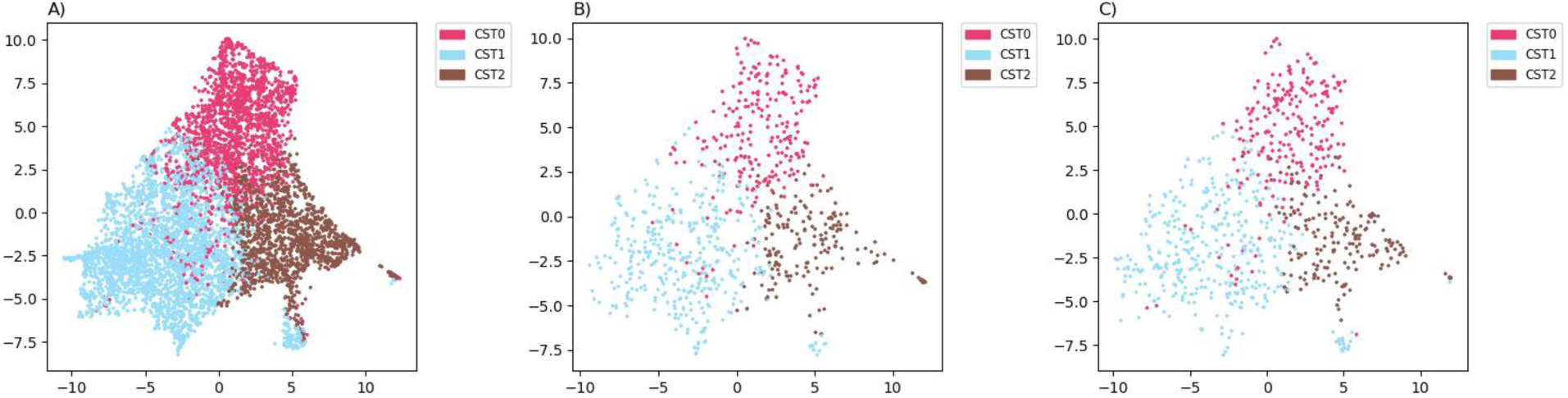
PACMAPs showing Manghi et al. [45] dataset with KMeans clustering labels (for three clusters: CST0, CST1, CST2), divided into A) the 80% training set, B) the 10% testing set, and C) the 10% validation set.

Labels assigned by StrataBionn were evaluated based on F1-score, recall, precision, and accuracy metrics for high and low training data availability scenarios. StrataBionn achieves 98.8-98.9% across all metrics with low training data availability, and 99% across all metrics in high training data availability scenarios (Figure 6A). We observed that this high level of accuracy was consistent across all label groups, with a slightly higher level of accuracy for less common CSTs given their size (Figure 6B). The least common CST, CST-2, was correctly identified in all 184 samples which truly belonged to that group, while only being mis-identified in two samples belonging to CST-1. CST-1, the most common subtype, is the one most commonly mis-assigned to a different cluster (7/369, 1.9%), likely due to the complicated boundaries between it and the other CSTs observed in figure 6B. StrataBionn is able to correctly identify these 98.1% of the time, indicating a strong ability to classify along complicated, non-linear boundaries.

**Figure 6.**
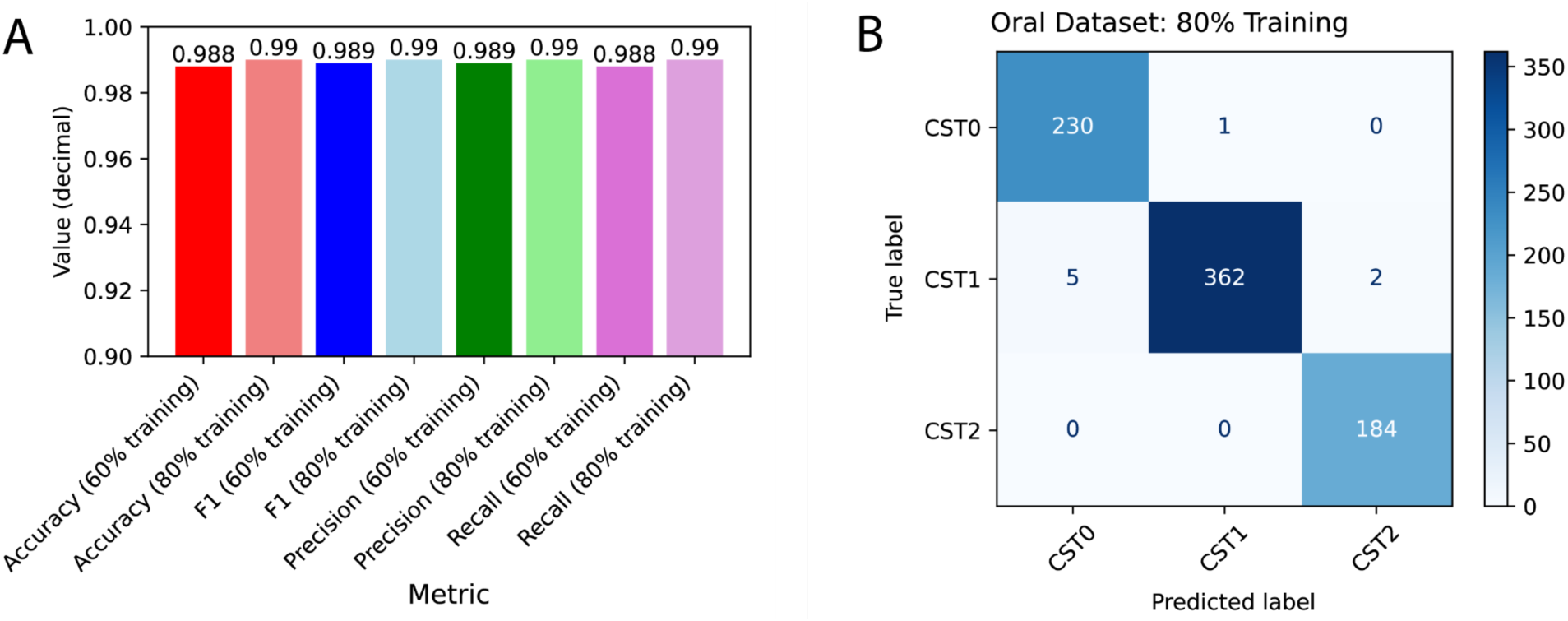
**(A)** Accuracy, F1 score, Precision, and Recall achieved by StrataBionn on the 60% validation and 80% validation clustered oral datasets. **(B)** Confusion matrix of StrataBionn classifications on the 80% classified oral data validation set. Counts on the diagonal correspond to successful assignments of communities by the methods to their true labels and the off-diagonals correspond to misassignments.

The in-house generated oral microbiome dataset containing 39 samples, and 47 additional samples from Baker et al [46]. were used to evaluate the ability of StrataBionn to apply classifications to novel non-vaginal microbiome datasets, and required no additional processing other than reformatting and column renaming. This new dataset was assigned labels using a classifier trained on the 80% training data availability training subset for the oral microbiome described above using only the bacteria species common to both datasets. All samples from Baker et al. [46] were classified as CST-1 by StrataBionn, matching the label of the samples with known CSTs with which they were compositionally most similar to (Fig. 7). In the in-house and Baker et al. [46] datasets, we identified 7 belonging to CST-0, and 79 samples in CST-1. No samples in these sets were identified as belonging to CST-2.

**Figure 7.**
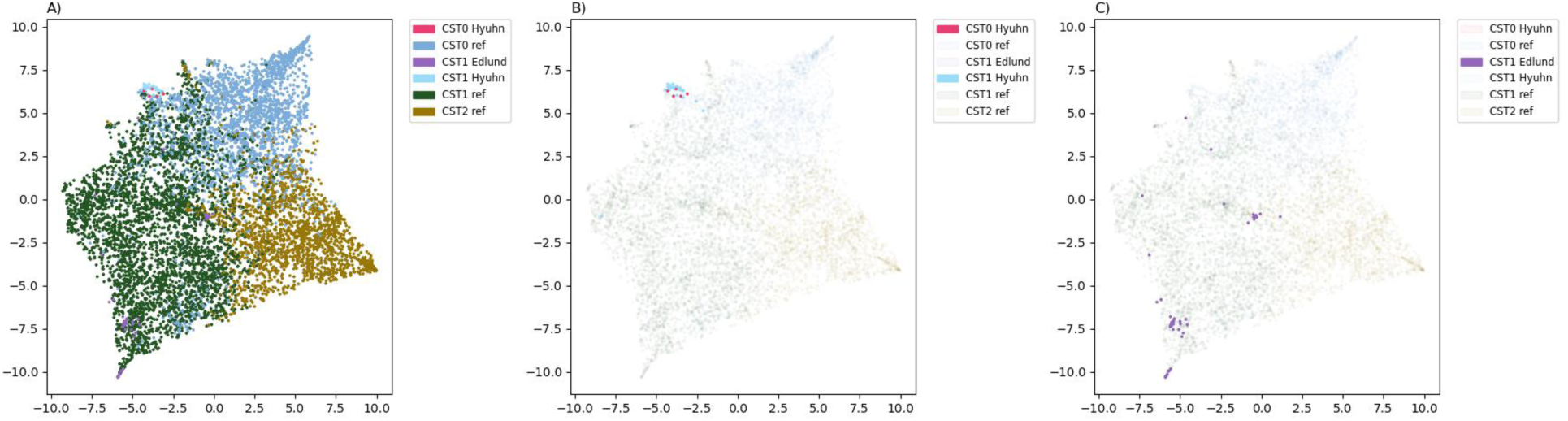
PACMAPs of Manghi et al. [45] oral microbiome reference dataset classifications, Baker et al. [46] oral microbiome dataset classification, and in-house oral microbiome dataset classifications generated by StrataBionn. In figure (A), both datasets are displayed with equal opacity. In figure (B), classified in-house generated data is displayed at full opacity while reference data is displayed at 5% opacity. In figure (C), Baker et al. [46] oral microbiome data is displayed at full opacity while reference data is displayed at 5% opacity.

## Discussion

There is increasing evidence of the implication of community-level microbial shifts with the increased likelihood of disease, and human health more broadly [19,55–66]. This puts an increased importance on our ability to accurately infer the microbial communities and discriminate those that are commonly associated with healthy outcomes from those commonly associated with disease. The complexity of microbial community data, characterized by high dimensionality, compositional sparsity, and technical noise, requires robust, interpretable, and transferable classification frameworks. The classification process is generally abstracted away and hidden from the user, concealing useful information regarding how assignments are determined during the classification process which hinders the interpretability of those classifications. Additionally, current classification methods are fundamentally constrained to a single dataset with a fixed set of bacteria species, preventing meaningful classifications derived from one study from being applied generally to additional studies for the same microbiome. To address these issues, we developed StrataBionn, an Artificial Neural Network (ANN)-based tool designed to overcome the limitations of extant classifiers by providing superior accuracy and cross-study applicability. By integrating transparent visualization and perturbation analysis, StrataBionn transitions microbiome “state-typing” from a “black-box” abstraction to an interpretable diagnostic process, where researchers can investigate the compositional causality of the inferred community labels in a community. Although StrataBionn has been developed for microbial communities, it could likely be adapted to study any ecological community data for which abundant samples and training are available due to its community-independent composition based classification architecture.

### Comparative Performance and Architecture Selection

Our results demonstrate that StrataBionn consistently outperforms both the k-nearest neighbor-based approach of VALENCIA and a traditional Random Forest (RF) classifier implemented in our package. While RF models showed competitive performance when compared to k-nearest neighbor approaches, the ANN architecture was selected for the basis of StrataBionn due to its superior computational efficiency and accuracy at scale. Notably, StrataBionn achieved a **12-13% increase** in F1-score and precision over VALENCIA when benchmarked against the vaginal microbiome, a dataset and type of community that VALENCIA was specifically designed to resolve. The performance gain was most pronounced in the classification of sub-CSTs (e.g., CSTs IV-A and IV-C0), which often present overlapping composition profiles that confound less sensitive models. Compared to the Random Forest classifier, StrataBionn achieves an approximately 2% increase across all the same metrics, with the difference being slightly more pronounced with more training data. The performance metric differences between the 60% training dataset (7,935 training samples) and the 80% training dataset (10,580 training samples) were negligible in our testing, but the 80% training dataset did achieve slightly higher performance. More specifically, we can attribute this increase in correct assignments to a higher precision for all sub-CSTs of CST I and III, and higher precision for CSTs IV-A, IV-C0, and V (Fig. 3). This is important because often CST-IVs are associated with increased likelihood of disease, and thus our method can dramatically reduce false positives in disease susceptibility screenings by correctly attributing samples to their true, non-CST-IV type, without degraded recall ability. What is less known is how the observed differences among CST-IV subgroups are differentially associated with disease, making the distinction of those communities relevant for any future study aimed at discovering the compositional and causal contributions of these community differences to disease.

The marginal performance increase observed when expanding the training set from 60% to 80% suggests that StrataBionn reaches an early accuracy plateau, indicating high data efficiency. This is particularly important given that many datasets are going to be sample limited, and in those circumstances, the strategy for the partition of the samples for training, testing and validation might differ. While France et al. [37] argued that VALENCIA’s assignments may better reflect biological states than the hierarchical clustering (HC) labels, our objective was to validate the learning fidelity of the model. By using HC labels as a ground-truth proxy, we demonstrated that StrataBionn possesses a superior capacity to internalize and replicate complex, user-defined classification logic. The ANN learns classification assignments based on their compositions during an iterative training process, where training is continued until accuracy improvements reach a maximum. The result is a high accuracy, deterministic trained model which can apply consistent classifications to novel data. As the training process learns distributions based on user-provided labels, it is not inherently tied to any one microbiome, which has been a major limitation of previous models. Additionally, by allowing the user to specify a set of bacteria species to consider during this process, disregarding other species, StrataBionn can produce additional trained models which can be applied across separate datasets for meta-analysis.

In the PACMAP visualization of the data, it can be seen that some samples assigned to CSTs appear isolated from the general clusters. This visualization can help researchers identify samples with low probability of assignment. In Supplementary Table 1, we provide the probabilities of assignment for a subset of samples from the vaginal microbiome that seem isolated from their clusters. In these samples, the general trend is that they share a low probability of assignment to their clusters. Because our method provides an empirical probability of assignment, researchers can perform a post-assignment filter to declare ambiguities.

### Interpretability and Compositional Causality

A persistent critique of deep learning in biology is the lack of model transparency. We took a novel approach to address this concern by including a classification boundary visualization tool and a perturbation analysis utility, which increases transparency in the classification process. More specifically, StrataBionn addresses the issue of interpretability via two distinct mechanisms:

1. **Feature-Space Visualization:** By projecting classification boundaries onto two-dimensional bacterial abundance coordinate planes, researchers can visually verify the “decision logic” of the model. By allowing the user to select two bacteria for the x and y axis and displaying the classification assignment in a scatter plot, insight as to what bacteria correspond to which CST can be attained. For example, we can inspect which bacteria are correlated with the different sub-CSTs by searching for a bacteria species where all samples appear to vary widely in abundance, appearing distributed across an axis. Then, we can find another species varying widely in abundance and observe the clustering of certain CSTs, displayed in different colors, to determine the features and thresholds which lead to their assignment. Examples of these plots and their interpretations are provided in supplementary figures 5-7.
2. **Stratified Perturbation Analysis:** Stratabionn’s perturbation analysis utility quantifies the sensitivity of the model to specific taxa. By permuting the counts of individual species and measuring the resulting decay in F1-score, StrataBionn identifies which bacteria are computationally “essential” for a specific CST assignment. StrataBionn provides the impact of individual species perturbations for F1, precision, recall, and accuracy metrics, providing additional insight into the impact of single species into the increased sensitivity or specificity of the assignment to communities (supplementary figure 8).

We propose that users implement the classification algorithm and use the tools provided by StrataBionn to further investigate the compositional causality of specific community states. We anticipate that StrataBionn will enable researchers to identify emerging properties that cause microbial communities to differ across host populations in datasets of increasing sample size and global distribution.

### Generalizability and Metadata Integration

Unlike previous models fundamentally constrained to fixed taxa sets or specific datasets, StrataBionn’s architecture is microbiome-agnostic. The ability to specify subsets of bacteria for model training facilitates meta-analyses across disparate studies where sequencing depths or taxonomic resolutions may vary. This flexibility is critical as the field moves toward large-scale longitudinal cohorts involving the gut-brain axis, cardiovascular health, and susceptibility to sexually transmitted infections (STIs).

The growing body of evidence confirming the mechanistic connections between the microbiome and human health highlights a clear and increasing demand for highly accurate analytical tools. These tools must also be adaptable to the study of the diverse microbial communities associated with various hosts. Accurate characterization of the microbiome is essential to gain insights into the gut-brain axis in neurodegenerative and psychological disorders [19,55], understanding the role of various microbiotas in cardiovascular health [56,63,67], and determining susceptibility to respiratory, renal, and sexually transmitted infections [57–64]. Within the gut microbiome, a more granular understanding of the correlations between Community State Types (CSTs) and pathology could enable targeted therapeutic interventions. These include fecal microbiota transplants (FMT) designed to engraft disease-resistant profiles or the use of pre- and probiotics to modulate the microbiome toward a protective state [11,19,65,66]. Indeed, CST-level analyses have already proven more successful than lower-resolution studies in establishing the correlations necessary to identify and treat conditions such as Celiac disease [65,66]. However, because the performance of a state-typing model directly impacts the diagnostic accuracy of microbiome-disease associations, algorithms must provide both maximal confidence and biological explainability. StrataBionn addresses these requirements by delivering high-speed, consistent, and interpretable labeling, allowing researchers to interrogate the classification process without compromising predictive accuracy.

### Technical Considerations and Limitations

StrataBionn was pre-configured with default hyperparameter values which allow the network to accurately fit most classifications. Should the user desire to do so, they are enabled to manually tune the hyperparameters to maximize the classification quality. We distribute the tool in a GitHub repository that can be easily accessed by users and contains a provided conda environment to quickly get started. In most cases, the user needs only to activate the environment and provide labeled training and testing data to create a working classifier.

Despite the advantages of ANN architectures, the risk of overfitting remains a primary concern. Neural network based classifiers inherit a concern of overfitting, where instead of learning patterns in data, the model learns to memorize the samples in a dataset. We mitigated this by implementing an “early stopping” protocol, where training is terminated once loss for an independent test set plateaus, or decreases for several epochs. In addition to validation based early stopping, the network depth was purposefully constrained to mitigate the risk of overfitting to, or “memorizing” noise inherent to small-to-medium-sized datasets. If overfitting occurs, this parameter is tunable and users can modify them accordingly. One of the benefits of our ANN, as implemented in StrataBionn, is its ability to correctly assign community types in the presence of non-linear boundaries. This is more clearly shown in the case of the oral microbiome (Figure 6B). These types of boundary classifications,which are likely frequent in microbiome data, are problematic with simpler approaches like the nearest centroid classification algorithms.

Classifications from a pre-existing study with known biologically-relevant and composition-based CST labels [29] were used as a reference when validating StrataBionn’s performance on the vaginal microbiome. As there are no generally accepted reference classifications for the oral microbiome, we employed a K-means clustering model with 3 clusters (see supplementary figures 3 and 4) to create naive reference classifications to validate our model on oral microbiota data. As StrataBionn is designed to learn sample composition patterns from training data with curated labels, the biological meaning behind the generated labels for the oral microbiome are unimportant to the validation of the tool.

Studies have shown that the composition of microbiomes are inherently dynamic, changing due to environmental and internal factors [68–70]. StrataBionn is limited by its inability to consider temporal data, which could enhance the labelling ability of microbiome classification methods. Future work could incorporate this data to gain a greater insight into the microbiome of an individual, and how these state-change events correlate to disease acquisition and susceptibility. Future iterations could incorporate **Recurrent Neural Networks (RNNs)**, **Long Short-Term Memory (LSTM)** networks, or **Transformers** to process temporal longitudinal data. Such an evolution would allow for the prediction of state-change transitions, potentially identifying early warning signs of dysbiosis before clinical symptoms manifest. As this information is collected from the compilations of data points taken at discrete times, the high accuracy of StrataBionn allows for the collection and aggregation of this data to further investigate the temporal evolution of microbiome state-type in individuals.

As the field of microbiome research rapidly progresses, demanding faster and more accurate community state typing, we attempt to contribute a solution. With StrataBionn, we offer a robust and scalable classification tool designed to accommodate diverse studies, including the analysis of ecological community data. Our goal is to enable future research to thoroughly investigate the impact of microfauna composition on human and possibly ecosystem health. StrataBionn achieves this by accurately classifying samples and providing tools to explore the decision boundaries that govern the classification process.

## Availability of data and materials

Information on how to obtain the vaginal microbiome datasets from France et al. [37] and Hickey et al. [44], and the oral microbiome datasets from Manghi et al. [45] and Baker et al. [46] can be found in their respective papers (we used the relative abundances reported by each one of these studies). Nevertheless, a freeze of the tables with the relative abundances from these studies is available in our github in the data folder. StrataBionn is available at https://github.com/KelleyCornejoLabs/Microbiome_Classification. The raw FastQC for the in-house generated oral microbiome dataset is available at NCBI-SRA under BioProject PRJNA1445365.

## Supplementary Materials

**Supplementary Table 1.** This table shows the probabilities of assignment to each Sub-CST for all samples which appeared isolated compared to surrounding samples. Isolated in this context was defined as samples whose closest neighbor’s euclidean distance was above a threshold, after undergoing a PACMAP transformation. 13 such samples were discovered, and their assignment confidence values were gathered in the table above. We can see that samples found to be distant from other samples after a PACMAP transform, which groups samples of a similar composition, tend to receive a relatively low reported confidence value from StrataBionn.

| sampleID | subCST | Pct I-A | Pct I-B | Pct II | Pct III-A | Pct III-B | Pct IV-A | Pct IV-B | Pct IV-C0 | Pct IV-C1 | Pct IV-C2 | Pct IV-C3 | Pct IV-C4 | Pct V |
| --- | --- | --- | --- | --- | --- | --- | --- | --- | --- | --- | --- | --- | --- | --- |
| X11601Vag1 | IV-C0 | 0.18424 | 0.21925 | 0.02700 | 0.00000 | 0.00000 | 0.03049 | 0.10513 | **0.37480** | 0.05866 | 0.00018 | 0.00003 | 0.00018 | 0.00002 |
| X13206Vul1 | IV-C0 | 0.16653 | 0.18997 | 0.00887 | 0.00000 | 0.01007 | 0.06226 | 0.06902 | **0.46464** | 0.02541 | 0.00127 | 0.00028 | 0.00139 | 0.00028 |
| 1476 | IV-B | 0.00001 | 0.00000 | 0.00020 | 0.00000 | 0.00000 | 0.01215 | **0.98763** | 0.00001 | 0.00000 | 0.00000 | 0.00000 | 0.00000 | 0.00000 |
| 3518 | IV-C3 | 0.00175 | 0.02149 | 0.22428 | 0.00000 | 0.00001 | 0.00000 | 0.00016 | 0.03174 | 0.20135 | 0.00014 | **0.51753** | 0.00154 | 0.00001 |
| 3689 | III-B | 0.00702 | 0.02168 | 0.00523 | 0.09092 | **0.86839** | 0.00012 | 0.00057 | 0.00196 | 0.00399 | 0.00002 | 0.00004 | 0.00005 | 0.00002 |
| 5541 | IV-C1 | 0.32912 | 0.01465 | 0.03246 | 0.00000 | 0.00000 | 0.00000 | 0.00006 | 0.04021 | **0.58345** | 0.00001 | 0.00004 | 0.00000 | 0.00000 |
| 6497 | I-A | **0.77430** | 0.21516 | 0.00852 | 0.00004 | 0.00015 | 0.00000 | 0.00000 | 0.00136 | 0.00008 | 0.00002 | 0.00001 | 0.00001 | 0.00035 |
| 8440 | I-A | **0.98149** | 0.01078 | 0.00773 | 0.00000 | 0.00000 | 0.00000 | 0.00000 | 0.00000 | 0.00000 | 0.00000 | 0.00000 | 0.00000 | 0.00000 |
| 9348 | IV-C0 | 0.15929 | 0.13283 | 0.06770 | 0.00000 | 0.00002 | 0.00000 | 0.00001 | **0.35498** | 0.28195 | 0.00242 | 0.00043 | 0.00037 | 0.00000 |
| 10640 | III-A | 0.00944 | 0.09548 | 0.01248 | **0.87770** | 0.00436 | 0.00000 | 0.00000 | 0.00039 | 0.00015 | 0.00000 | 0.00000 | 0.00000 | 0.00000 |
| 11537 | I-A | **0.58060** | 0.10947 | 0.26313 | 0.00000 | 0.00000 | 0.00000 | 0.00000 | 0.02705 | 0.01963 | 0.00001 | 0.00011 | 0.00001 | 0.00000 |
| 11829 | I-A | **0.80962** | 0.17966 | 0.01056 | 0.00000 | 0.00000 | 0.00000 | 0.00008 | 0.00002 | 0.00001 | 0.00001 | 0.00000 | 0.00000 | 0.00005 |
| 12807 | III-B | 0.00879 | 0.02412 | 0.00297 | 0.00750 | **0.88996** | 0.00043 | 0.00570 | 0.01193 | 0.04712 | 0.00023 | 0.00028 | 0.00092 | 0.00007 |

**Supplementary Figure 1.**
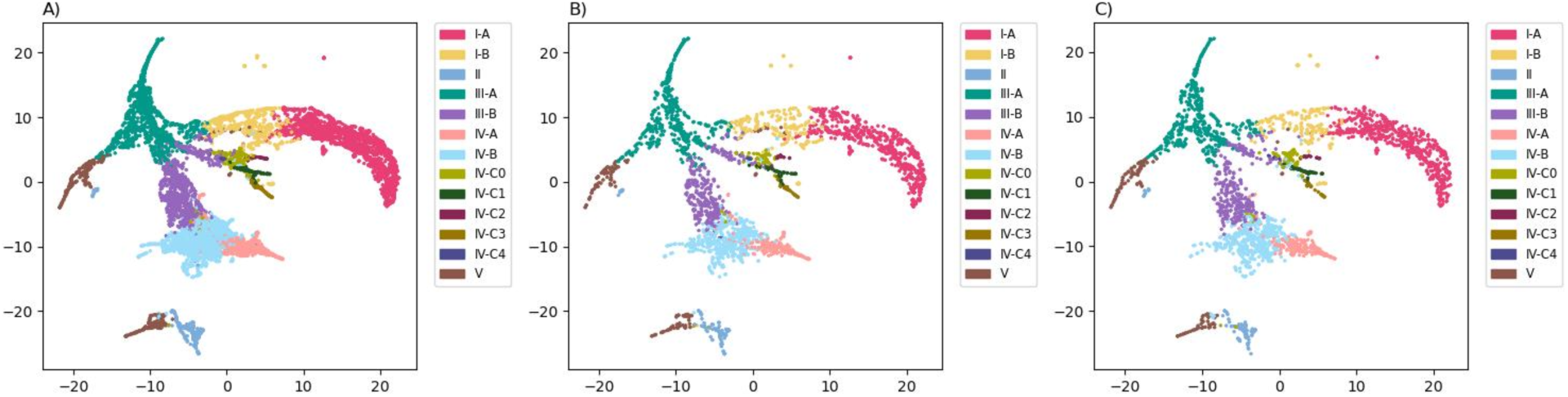
PACMAPs showing classifications from France et al. for A) the 60% training set, B) the 20% testing set, and C) the 20% vaginal microbiome validation set.

**Supplementary Figure 2.**
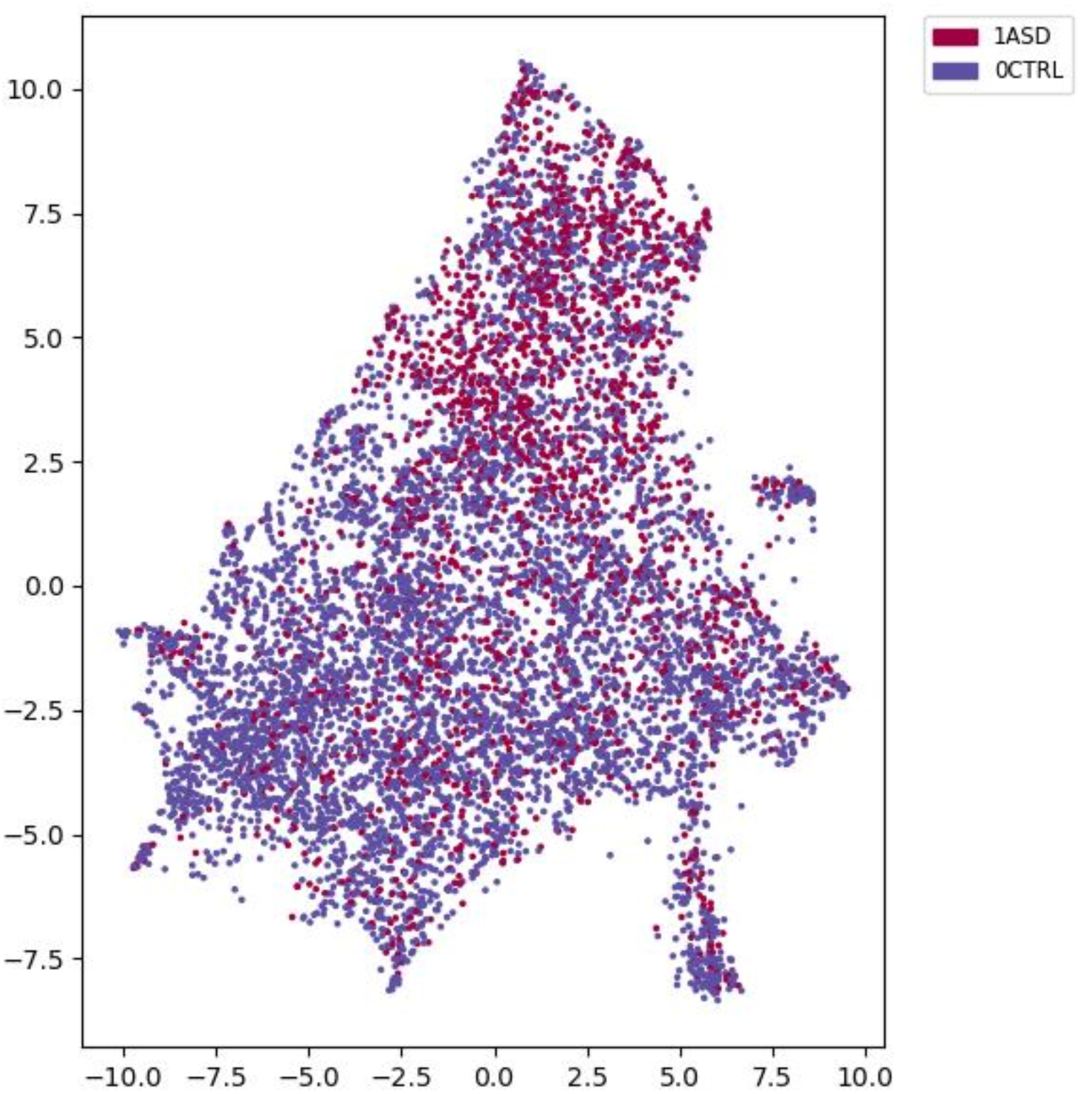
PACMAP containing 7,812 samples from Manghi et al. with Autism Spectrum Disorder (ASD) and control labels. While samples labeled ASD occur more commonly near the top of the PACMAP, ASD and control labeled samples co-occur throughout the entire figure. The lack of distinct clusters implies this classification scheme is not rooted in sample composition.

**Supplementary Figure 3.**
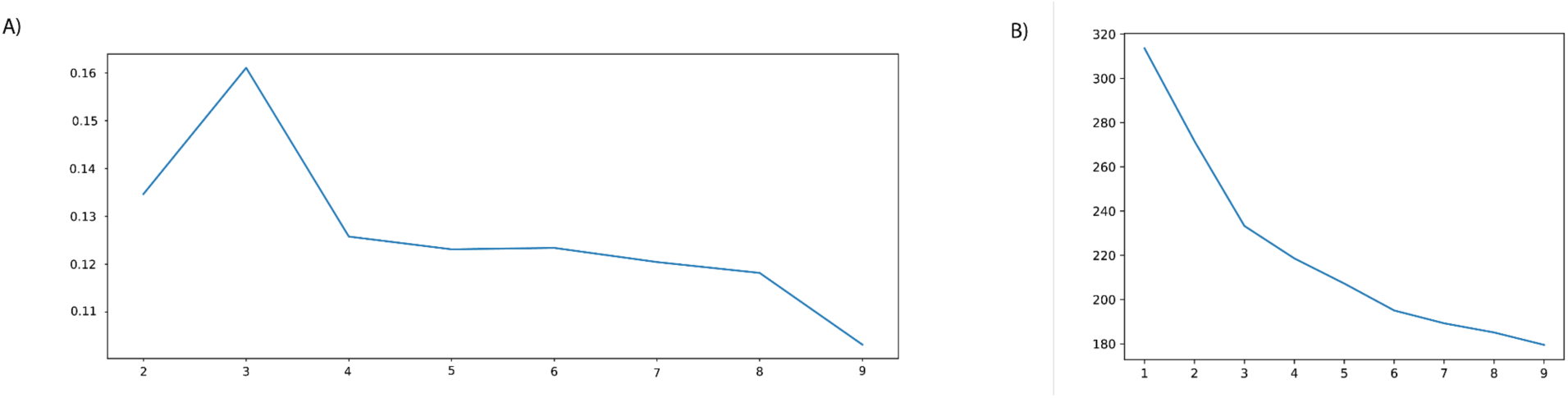
A) Graph displaying silhouette scores for k clusters for naive classification of oral microbiome data using K-means clustering. We observe the silhouette score is highest using 3 clusters. B) Elbow plot showing within-cluster sum of squares (inertia) against number of clusters. Here we can identify that the inertia begins to slow when the cluster count reaches 3, which supports our decision to use three oral CST labels.

**Supplementary Figure 4.**
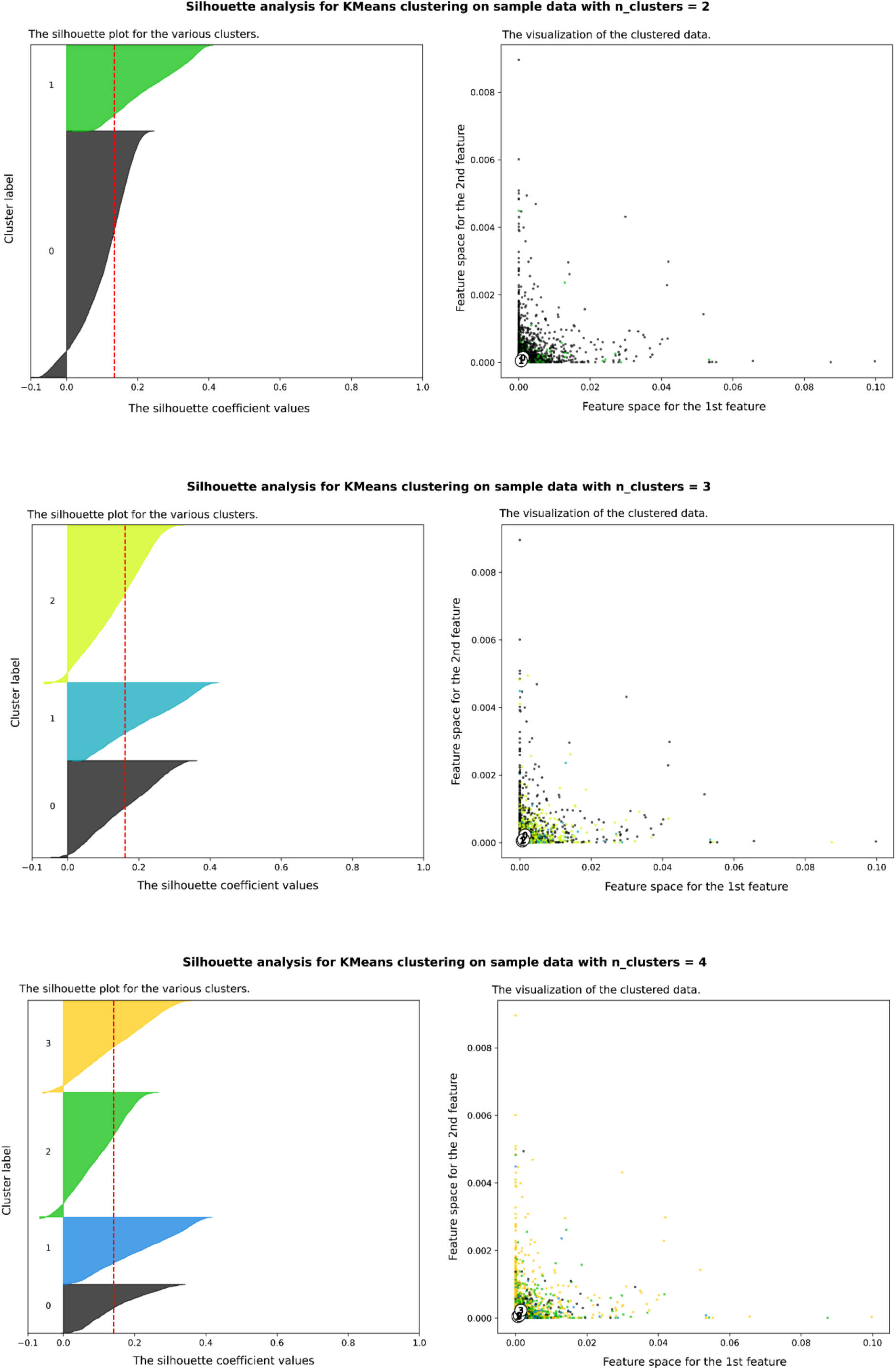
Silhouette analysis plots showing the silhouette score for each sample in the oral microbiome dataset. Silhouette scores were calculated using naive K-means clustering-derived labels, with two (A), three (B), and four (C) clusters being tested. Using three clusters yields the highest average silhouette score, indicating the highest cluster separation.

**Supplementary Figure 5.**
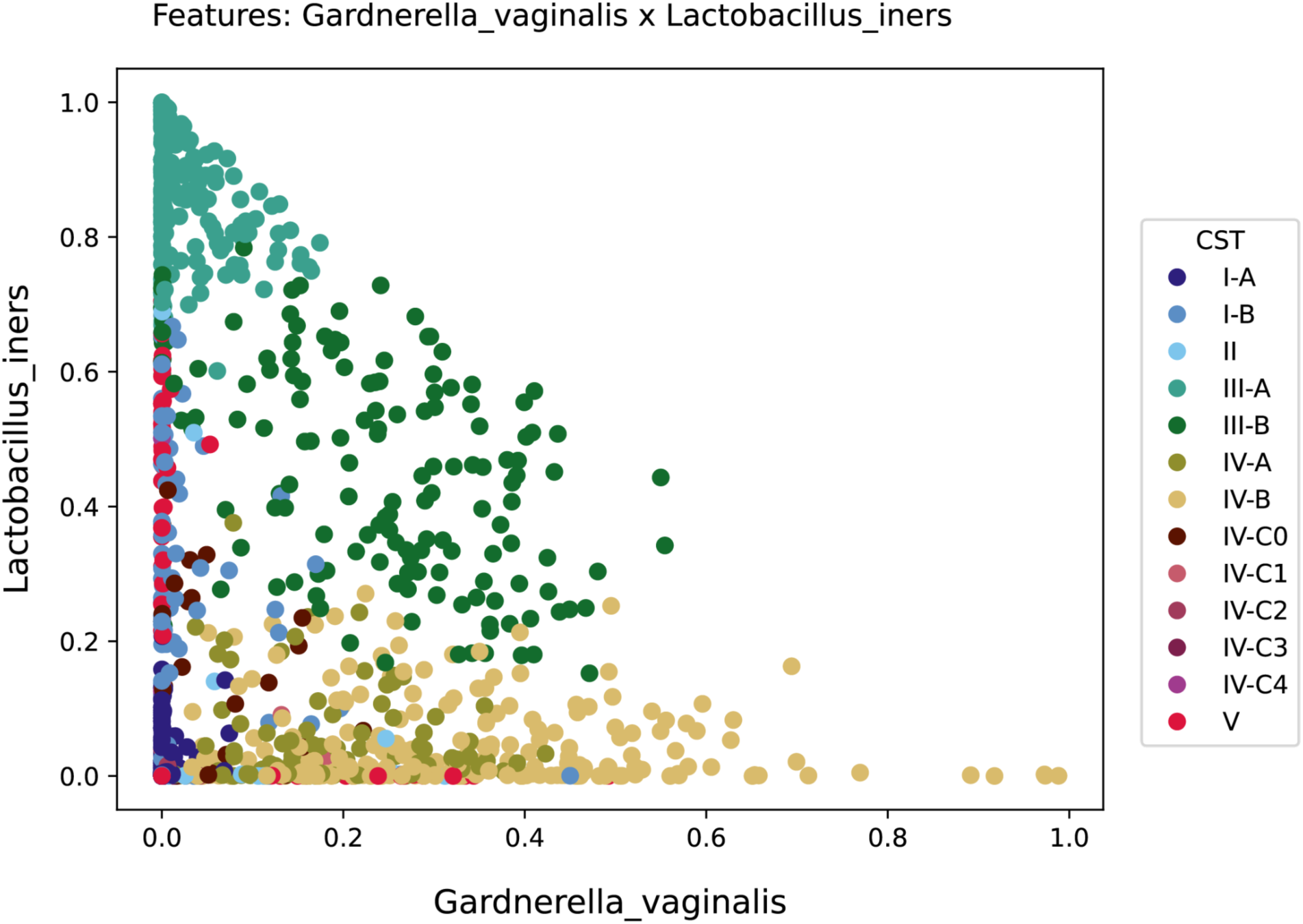
This figure shows the decision boundaries learned by StrataBionn to classify the vaginal microbiome, focusing on the bacteria species *Gardnerella vaginalis* and *Lactobacillus iners*. We can clearly see in this graph that samples dominated by *G. vaginalis* (>60% composition) are always assigned to CST IV-B. Interestingly while there are many CST IV-B samples with a high proportion of *G. vaginalis*, few of them also contain as much *L. iners* as can be found in CST IV-B samples containing less *G. vaginallis*, although it would be possible.

**Supplementary Figure 6.**
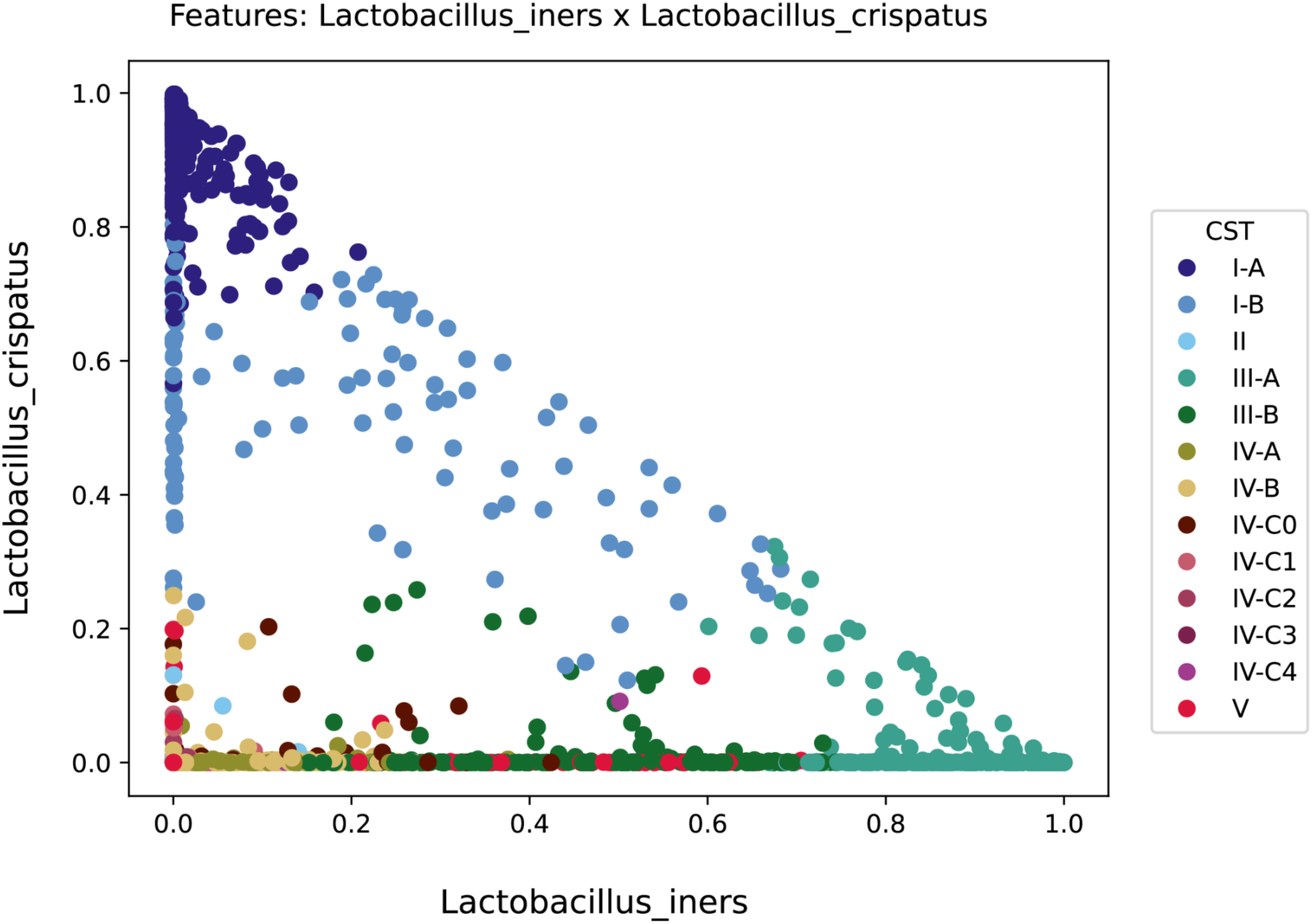
This figure shows the decision boundaries learned by StrataBionn to classify the vaginal microbiome, focusing on the bacteria species *Lactobacillus crispatus* and *Lactobacillus iners*. We can observe that CSTs I-A and I-B have a non-linear boundary when a sample is composed of ∼70% *L. crispatus*. While CST I-A generally clusters in the corner where both bacteria species are low in relative abundance, this plot shows that the majority of the composition of these samples is *L. crispatus*. CST I-B has a complex but clear decision boundary with other CSTs. It is commonly assigned when a sample contains >∼25% *L. crispatus*, but can occur with lower proportions of *L. crispatus* near ∼50% L. iners. Sub-CSTs III-A and III-B seem to have a fuzzy decision boundary when a sample contains ∼75% *L. iners*. Samples containing >∼30% *L. crispatus* are generally assigned to neither III-A nor III-B, and are exclusively assigned to sub-CSTs of CST I. We also observe the compositional similarities between Sub-CSTs of CST I and CST III, as both share long boundaries where a single, clear line is difficult to draw using these axes.

**Supplementary Figure 7.**
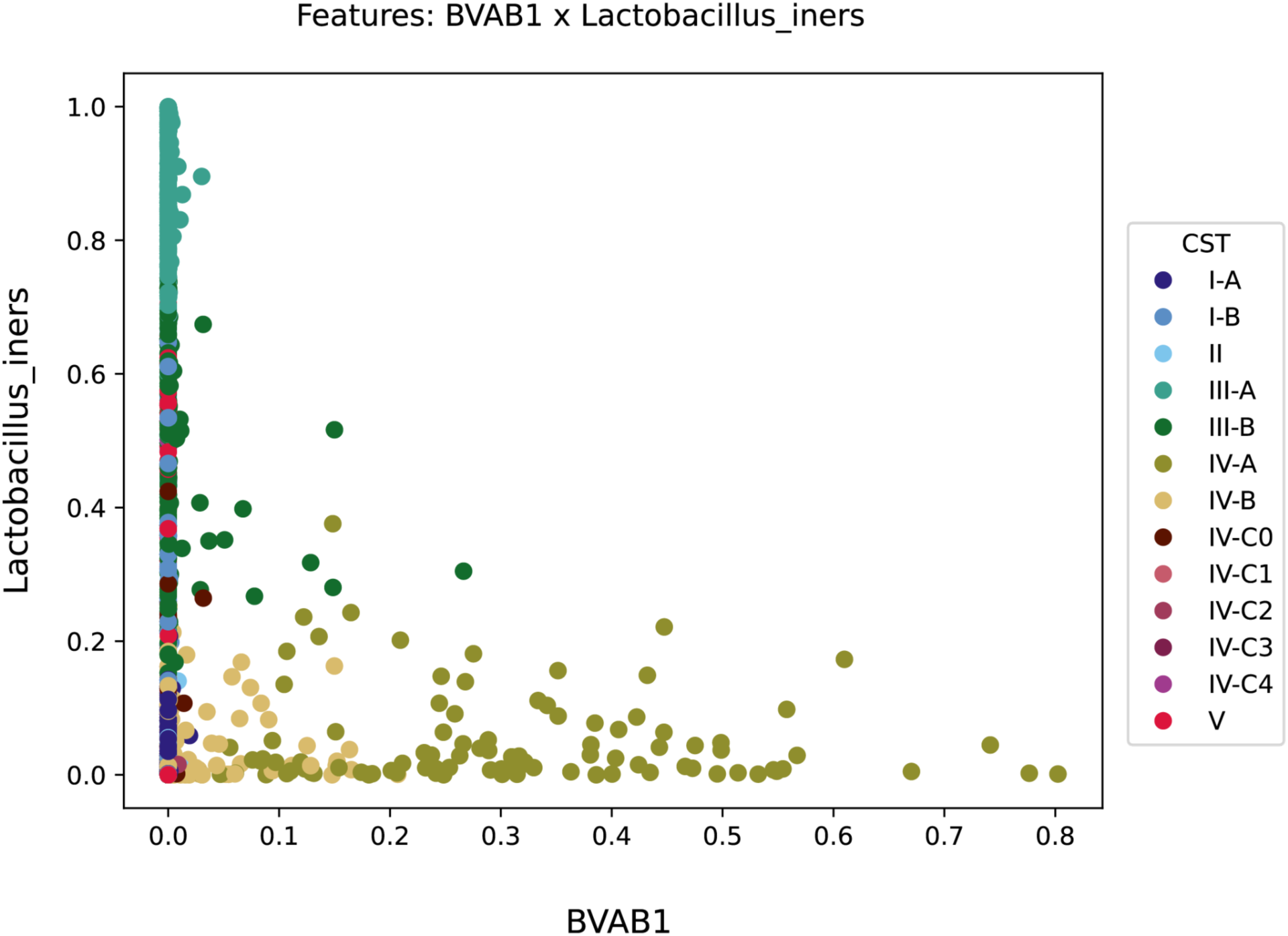
This figure shows the decision boundaries learned by StrataBionn to classify the vaginal microbiome, focusing on the bacteria species “*Candidatus Lachnocurva vaginae*” (formerly BVAB1) and *Lactobacillus iners*. In this plot we can observe that while most CSTs have relatively low abundances of “*Ca. Lachnocurva vaginae*”, Sub-CSTs IV-A, IV-B, and III-B are exceptions. Such samples with high levels of *L. iners* are generally classified as Sub-CST III-A, and samples with >∼30% “*Ca*. *Lachnocurva vaginae*” are always classified as Sub-CST IV-A.

**Supplementary Figure 8.**
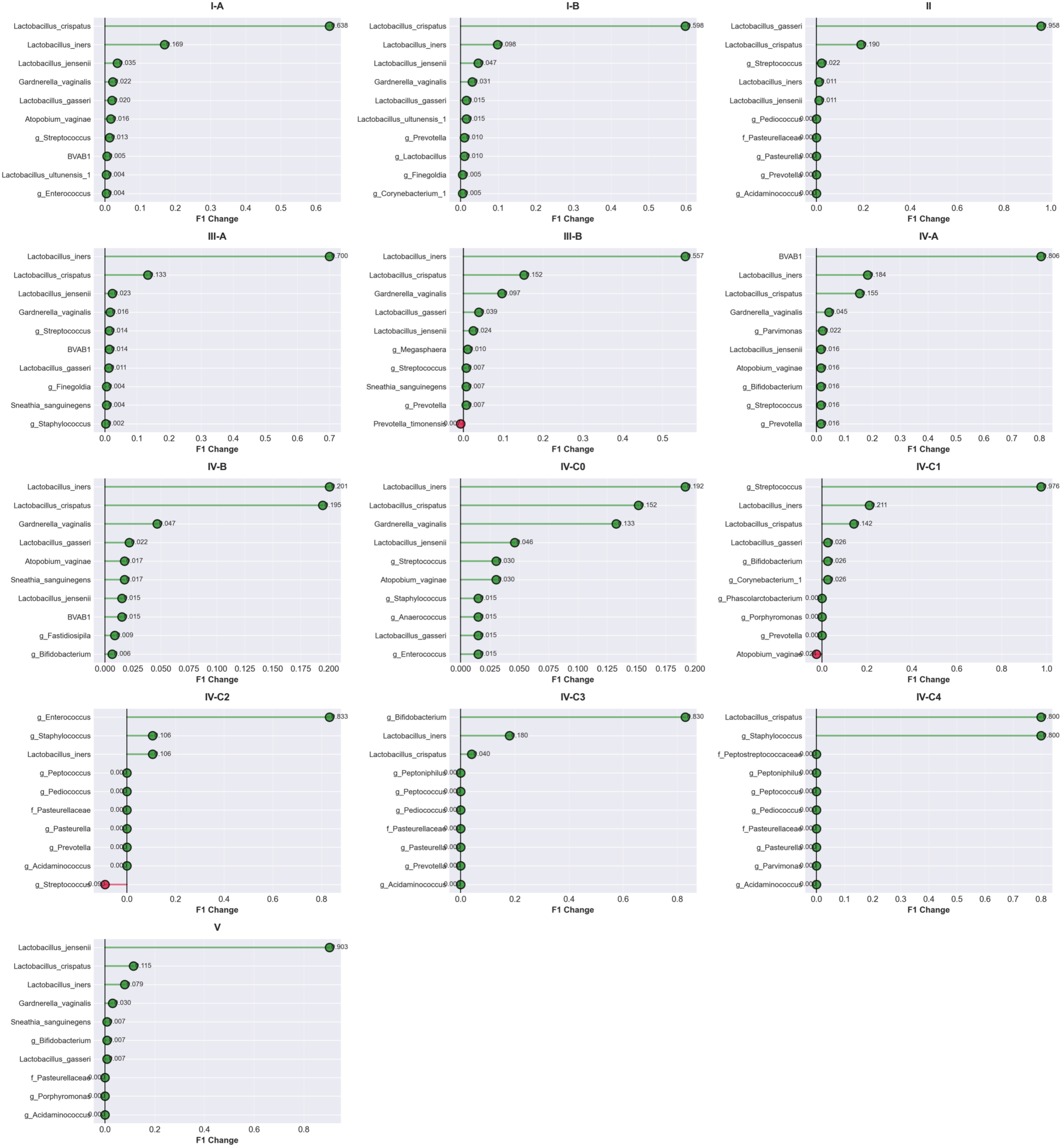
Cleveland plots showing the impact of different species perturbation on the F1 score assignment to each CST type. We propose the use of this visualization in combination with the Feature space visualization from supplementary Figures 5-7 to identify defining features (species) that are relevant for the assignment of community labels.

**Supplementary Figure 9.**
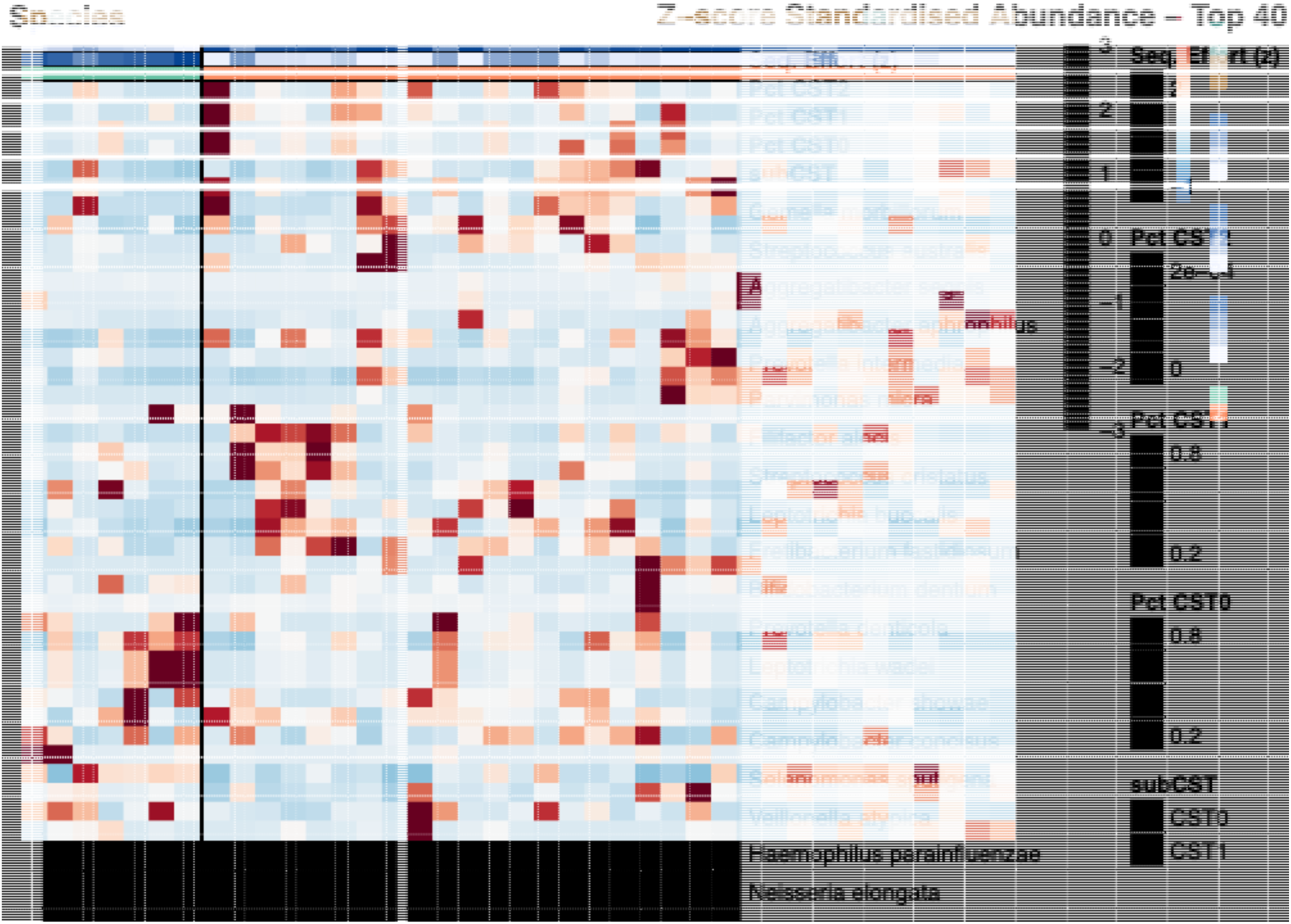
Heatmap for top 50 species in the oral dataset sequenced for this study. The values in the cells correspond to z normalized values of relative abundance and the annotation in the top corresponds to the subCST type that each sample was assigned to (CST0, CST1) and the probabilities of assignment to each CST type.

In Supplementary Table 2. It can be seen that the uncertainty in the assignment, estimated as more evenness (Shannon-information index) in the values of probability of assignment is not significantly correlated with sequencing effort. Shannon diversity index in the probability of assignment was estimated as

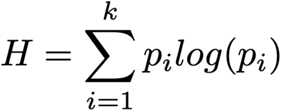

A normalized version of Shannon was estimated as

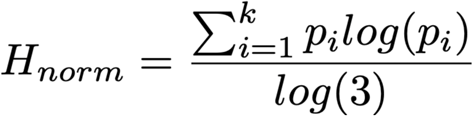

Comparisons were made between both the standard Shannon diversity index and the normalized version of it.

**Supplementary Table 2.** Correlation values and probability.

Correlation Matrix with P-values
| Row | Column | Corr | P |
| --- | --- | --- | --- |
| Sequencing_effort | H_norm | -0.31 | 0.0581 |
| Sequencing_effort | H | -0.31 | 0.0581 |
| H_norm | H | 1.00 | 0.0000 |

## References

1. Björk JR, Bolte LA, Maltez Thomas A, Lee KA, Rossi N, Wind TT, et al. Longitudinal gut microbiome changes in immune checkpoint blockade-treated advanced melanoma. Nat Med. Nature Publishing Group; 2024;30:785–96. 10.1038/s41591-024-02803-3

2. Clauss M, Gérard P, Mosca A, Leclerc M. Interplay Between Exercise and Gut Microbiome in the Context of Human Health and Performance. Front Nutr [Internet]. Frontiers; 2021 [cited 2026 Jan 7];8. 10.3389/fnut.2021.637010

3. Fan Y, Pedersen O. Gut microbiota in human metabolic health and disease. Nat Rev Microbiol. Nature Publishing Group; 2021;19:55–71. 10.1038/s41579-020-0433-9

4. Gunjur A, Shao Y, Rozday T, Klein O, Mu A, Haak BW, et al. A gut microbial signature for combination immune checkpoint blockade across cancer types. Nat Med. Nature Publishing Group; 2024;30:797–809. 10.1038/s41591-024-02823-z

5. Jewell MD, van Moorsel SJ, Bell G. Presence of microbiome decreases fitness and modifies phenotype in the aquatic plant Lemna minor. AoB PLANTS. 2023;15:plad026. 10.1093/aobpla/plad026

6. Kandalai S, Li H, Zhang N, Peng H, Zheng Q. The human microbiome and cancer: a diagnostic and therapeutic perspective. Cancer Biol Ther. 24:2240084. 10.1080/15384047.2023.2240084

7. Liu B-N, Liu X-T, Liang Z-H, Wang J-H. Gut microbiota in obesity. World J Gastroenterol. 2021;27:3837–50. 10.3748/wjg.v27.i25.3837

8. Gould AL, Zhang V, Lamberti L, Jones EW, Obadia B, Korasidis N, et al. Microbiome interactions shape host fitness. Proc Natl Acad Sci [Internet]. Proceedings of the National Academy of Sciences; 2018 [cited 2026 Jan 7]; 10.1073/pnas.1809349115

9. Ogunrinola GA, Oyewale JO, Oshamika OO, Olasehinde GI. The Human Microbiome and Its Impacts on Health. Int J Microbiol. 2020;2020:8045646. 10.1155/2020/8045646

10. Roelands J, Kuppen PJK, Ahmed EI, Mall R, Masoodi T, Singh P, et al. An integrated tumor, immune and microbiome atlas of colon cancer. Nat Med. Nature Publishing Group; 2023;29:1273–86. 10.1038/s41591-023-02324-5

11. Sasidharan Pillai S, Gagnon CA, Foster C, Ashraf AP. Exploring the Gut Microbiota: Key Insights Into Its Role in Obesity, Metabolic Syndrome, and Type 2 Diabetes. J Clin Endocrinol Metab. 2024;109:2709–19. 10.1210/clinem/dgae499

12. Varghese S, Rao S, Khattak A, Zamir F, Chaari A. Physical Exercise and the Gut Microbiome: A Bidirectional Relationship Influencing Health and Performance. Nutrients. 2024;16:3663. 10.3390/nu16213663

13. Xun W, Shao J, Shen Q, Zhang R. Rhizosphere microbiome: Functional compensatory assembly for plant fitness. Comput Struct Biotechnol J. 2021;19:5487–93. 10.1016/j.csbj.2021.09.035

14. Zhan M, Wang L, Xie C, Fu X, Zhang S, Wang A, et al. Succession of Gut Microbial Structure in Twin Giant Pandas During the Dietary Change Stage and Its Role in Polysaccharide Metabolism. Front Microbiol [Internet]. Frontiers; 2020 [cited 2026 Jan 7];11. 10.3389/fmicb.2020.551038

15. Durack J, Lynch SV. The gut microbiome: Relationships with disease and opportunities for therapy. J Exp Med. 2019;216:20–40. 10.1084/jem.20180448

16. Ghosh TS, Das M, Jeffery IB, O’Toole PW. Adjusting for age improves identification of gut microbiome alterations in multiple diseases. Turnbaugh P, Garrett WS, Lozupone CA, Turnbaugh P, editors. eLife. eLife Sciences Publications, Ltd; 2020;9:e50240. 10.7554/eLife.50240

17. Grosso F, Zanetti D, Sanna S. Causal relationships between gut microbiome and hundreds of age-related traits: evidence of a replicable effect on ApoM protein levels. Aging. 2025;17:1966–87. 10.18632/aging.206293

18. Gupta VK, Janda GS, Pump HK, Lele N, Cruz I, Cohen I, et al. Alterations in Gut Microbiome-Host Relationships After Immune Perturbation in Patients With Multiple Sclerosis. Neurol Neuroimmunol Neuroinflammation. Wolters Kluwer; 2025;12:e200355. 10.1212/NXI.0000000000200355

19. Hou K, Wu Z-X, Chen X-Y, Wang J-Q, Zhang D, Xiao C, et al. Microbiota in health and diseases. Signal Transduct Target Ther. Nature Publishing Group; 2022;7:135. 10.1038/s41392-022-00974-4

20. Lewis FMT, Bernstein KT, Aral SO. Vaginal Microbiome and Its Relationship to Behavior, Sexual Health, and Sexually Transmitted Diseases. Obstet Gynecol. 2017;129:643–54. 10.1097/AOG.0000000000001932

21. Madhogaria B, Bhowmik P, Kundu A. Correlation between human gut microbiome and diseases. Infect Med. 2022;1:180–91. 10.1016/j.imj.2022.08.004

22. Chen H, Jiang W. Application of high-throughput sequencing in understanding human oral microbiome related with health and disease. Front Microbiol [Internet]. Frontiers; 2014 [cited 2026 Feb 24];5. 10.3389/fmicb.2014.00508

23. Di Bella JM, Bao Y, Gloor GB, Burton JP, Reid G. High throughput sequencing methods and analysis for microbiome research. J Microbiol Methods. 2013;95:401–14. 10.1016/j.mimet.2013.08.011

24. Compositional analysis: a valid approach to analyze microbiome high-throughput sequencing data [Internet]. [cited 2026 Feb 24]. https://cdnsciencepub.com/doi/full/10.1139/cjm-2015-0821. Accessed 24 Feb 2026

25. Arumugam M, Raes J, Pelletier E, Le Paslier D, Yamada T, Mende DR, et al. Enterotypes of the human gut microbiome. Nature. Nature Publishing Group; 2011;473:174–80. 10.1038/nature09944

26. Costea PI, Hildebrand F, Arumugam M, Bäckhed F, Blaser MJ, Bushman FD, et al. Enterotypes in the landscape of gut microbial community composition. Nat Microbiol. 2018;3:8–16. 10.1038/s41564-017-0072-8

27. Callahan BJ, DiGiulio DB, Goltsman DSA, Sun CL, Costello EK, Jeganathan P, et al. Replication and refinement of a vaginal microbial signature of preterm birth in two racially distinct cohorts of US women. Proc Natl Acad Sci. Proceedings of the National Academy of Sciences; 2017;114:9966–71. 10.1073/pnas.1705899114

28. DiGiulio DB, Callahan BJ, McMurdie PJ, Costello EK, Lyell DJ, Robaczewska A, et al. Temporal and spatial variation of the human microbiota during pregnancy. Proc Natl Acad Sci. Proceedings of the National Academy of Sciences; 2015;112:11060–5. 10.1073/pnas.1502875112

29. Ravel J, Gajer P, Abdo Z, Schneider GM, Koenig SSK, McCulle SL, et al. Vaginal microbiome of reproductive-age women. Proc Natl Acad Sci U S A. 2011;108 Suppl 1:4680–7. 10.1073/pnas.1002611107

30. Ezugwu AE, Ikotun AM, Oyelade OO, Abualigah L, Agushaka JO, Eke CI, et al. A comprehensive survey of clustering algorithms: State-of-the-art machine learning applications, taxonomy, challenges, and future research prospects. Eng Appl Artif Intell. 2022;110:104743. 10.1016/j.engappai.2022.104743

31. Fang Y, Subedi S. Clustering microbiome data using mixtures of logistic normal multinomial models. Sci Rep. Nature Publishing Group; 2023;13:14758. 10.1038/s41598-023-41318-8

32. Gere A. Recommendations for validating hierarchical clustering in consumer sensory projects. Curr Res Food Sci. 2023;6:100522. 10.1016/j.crfs.2023.100522

33. Gloor GB, Macklaim JM, Pawlowsky-Glahn V, Egozcue JJ. Microbiome Datasets Are Compositional: And This Is Not Optional. Front Microbiol [Internet]. Frontiers; 2017 [cited 2026 Jan 7];8. 10.3389/fmicb.2017.02224

34. Liu Z, Yin X, Zhou Y, Li G, Chen K. Dissecting Microbial Community Structure and Heterogeneity via Multivariate Covariate-Adjusted Clustering [Internet]. arXiv; 2025 [cited 2026 Jan 7]. 10.48550/arXiv.2508.11036

35. Qiang J, Ding W, Kuijjer M, Quackenbush J, Chen P. Clustering Sparse Data With Feature Correlation With Application to Discover Subtypes in Cancer. IEEE Access Pract Innov Open Solut. 2020;8:67775–89. 10.1109/access.2020.2982569

36. Sinha R, Abu-Ali G, Vogtmann E, Fodor AA, Ren B, Amir A, et al. Assessment of variation in microbial community amplicon sequencing by the Microbiome Quality Control (MBQC) project consortium. Nat Biotechnol. 2017;35:1077–86. 10.1038/nbt.3981

37. France MT, Ma B, Gajer P, Brown S, Humphrys MS, Holm JB, et al. VALENCIA: a nearest centroid classification method for vaginal microbial communities based on composition. Microbiome. 2020;8:166. 10.1186/s40168-020-00934-6

38. Ronan T, Qi Z, Naegle KM. Avoiding common pitfalls when clustering biological data. Sci Signal. 2016;9:re6. 10.1126/scisignal.aad1932

39. Levner I. Feature selection and nearest centroid classification for protein mass spectrometry. BMC Bioinformatics. 2005;6:68. 10.1186/1471-2105-6-68

40. Ballabio D, Todeschini R. Multivariate Classification for Qualitative Analysis. Infrared Spectrosc Food Qual Anal Control. Academic Press; 2009. p. 83–100. 10.1016/B978-0-12-374136-3.00004-3

41. Blagus R, Lusa L. Class prediction for high-dimensional class-imbalanced data. BMC Bioinformatics. 2010;11:523. 10.1186/1471-2105-11-523

42. Sánchez Reyna AG, Mendoza-Gonzalez R, Luna-García H, Celaya Padilla JM, Morgan Benita JA, Espino-Salinas CH, et al. Synthetic data analysis for early detection of Alzheimer progression through machine learning algorithms. PeerJ Comput Sci. 2024;10:e2437. 10.7717/peerj-cs.2437

43. Ravel J, Gajer P, Abdo Z, Schneider GM, Koenig SSK, McCulle SL, et al. Vaginal microbiome of reproductive-age women. Proc Natl Acad Sci. Proceedings of the National Academy of Sciences; 2011;108:4680–7. 10.1073/pnas.1002611107

44. Hickey RJ, Zhou X, Settles ML, Erb J, Malone K, Hansmann MA, et al. Vaginal Microbiota of Adolescent Girls Prior to the Onset of Menarche Resemble Those of Reproductive-Age Women. mBio. American Society for Microbiology; 2015;6:10.1128/mbio.00097-15. 10.1128/mbio.00097-15

45. Manghi P, Filosi M, Zolfo M, Casten LG, Garcia-Valiente A, Mattevi S, et al. Large-scale metagenomic analysis of oral microbiomes reveals markers for autism spectrum disorders. Nat Commun. Nature Publishing Group; 2024;15:9743. 10.1038/s41467-024-53934-7

46. Baker JL, Morton JT, Dinis M, Alvarez R, Tran NC, Knight R, et al. Deep metagenomics examines the oral microbiome during dental caries, revealing novel taxa and co-occurrences with host molecules. Genome Res. Cold Spring Harbor Lab; 2021;31:64–74. 10.1101/gr.265645.120

47. FastQC [Internet]. 2015. https://qubeshub.org/resources/fastqc

48. Martin M. Cutadapt removes adapter sequences from high-throughput sequencing reads. EMBnet.journal. 2011;17:10–2. 10.14806/ej.17.1.200

49. Babraham Bioinformatics - Trim Galore! [Internet]. [cited 2026 Feb 24]. https://www.bioinformatics.babraham.ac.uk/projects/trim_galore/. Accessed 24 Feb 2026

50. Dobin A, Davis CA, Schlesinger F, Drenkow J, Zaleski C, Jha S, et al. STAR: ultrafast universal RNA-seq aligner. Bioinformatics. 2013;29:15–21. 10.1093/bioinformatics/bts635

51. Breitwieser FP, Baker DN, Salzberg SL. KrakenUniq: confident and fast metagenomics classification using unique k-mer counts. Genome Biol. 2018;19:198. 10.1186/s13059-018-1568-0

52. Pedregosa F, Varoquaux G, Gramfort A, Michel V, Thirion B, Grisel O, et al. Scikit-learn: Machine Learning in Python. J Mach Learn Res. 2011;12:2825–30.

53. Ansel J, Yang E, He H, Gimelshein N, Jain A, Voznesensky M, et al. PyTorch 2: Faster Machine Learning Through Dynamic Python Bytecode Transformation and Graph Compilation. 29th ACM Int Conf Archit Support Program Lang Oper Syst Vol 2 ASPLOS 24 [Internet]. ACM; 2024. 10.1145/3620665.3640366

54. Wang Y, Huang H, Rudin C, Shaposhnik Y. Understanding How Dimension Reduction Tools Work: An Empirical Approach to Deciphering t-SNE, UMAP, TriMAP, and PaCMAP for Data Visualization [Internet]. arXiv; 2021 [cited 2026 Feb 25]. 10.48550/arXiv.2012.04456

55. The Microbiota-Gut-Brain Axis | Physiological Reviews | American Physiological Society [Internet]. [cited 2026 Feb 24]. https://journals.physiology.org/doi/full/10.1152/physrev.00018.2018?rfr_dat=cr_pu. Accessed 24 Feb 2026

56. Sanchez-Rodriguez E, Egea-Zorrilla A, Plaza-Díaz J, Aragón-Vela J, Muñoz-Quezada S, Tercedor-Sánchez L, et al. The Gut Microbiota and Its Implication in the Development of Atherosclerosis and Related Cardiovascular Diseases. Nutrients. Multidisciplinary Digital Publishing Institute; 2020;12:605. 10.3390/nu12030605

57. Gosmann C, Anahtar MN, Handley SA, Farcasanu M, Abu-Ali G, Bowman BA, et al. Lactobacillus-Deficient Cervicovaginal Bacterial Communities Are Associated with Increased HIV Acquisition in Young South African Women. Immunity. 2017;46:29–37. 10.1016/j.immuni.2016.12.013

58. Brotman RM, Bradford LL, Conrad M, Gajer P, Ault K, Peralta L, et al. Association between Trichomonas vaginalis and vaginal bacterial community composition among reproductive-age women. Sex Transm Dis. 2012;39:807–12. 10.1097/OLQ.0b013e3182631c79

59. van Houdt R, Ma B, Bruisten SM, Speksnijder AGCL, Ravel J, de Vries HJC. Lactobacillus iners-dominated vaginal microbiota is associated with increased susceptibility to Chlamydia trachomatis infection in Dutch women: a case-control study. Sex Transm Infect. 2018;94:117–23. 10.1136/sextrans-2017-053133

60. Hilty M, Burke C, Pedro H, Cardenas P, Bush A, Bossley C, et al. Disordered Microbial Communities in Asthmatic Airways. PLOS ONE. Public Library of Science; 2010;5:e8578. 10.1371/journal.pone.0008578

61. Millares L, Ferrari R, Gallego M, Garcia-Nuñez M, Pérez-Brocal V, Espasa M, et al. Bronchial microbiome of severe COPD patients colonised by Pseudomonas aeruginosa. Eur J Clin Microbiol Infect Dis. 2014;33:1101–11. 10.1007/s10096-013-2044-0

62. Huang YJ, Nelson CE, Brodie EL, DeSantis TZ, Baek MS, Liu J, et al. Airway microbiota and bronchial hyperresponsiveness in patients with suboptimally controlled asthma. J Allergy Clin Immunol. Elsevier; 2011;127:372–381.e3. 10.1016/j.jaci.2010.10.048

63. Barros AF, Borges NA, Ferreira DC, Carmo FL, Rosado AS, Fouque D, et al. Is there Interaction Between Gut Microbial Profile and Cardiovascular Risk in Chronic Kidney Disease Patients? Future Microbiol. Taylor & Francis; 2015;10:517–26. 10.2217/fmb.14.140

64. Li L, Zhang Y-L, Liu X-Y, Meng X, Zhao R-Q, Ou L-L, et al. Periodontitis Exacerbates and Promotes the Progression of Chronic Kidney Disease Through Oral Flora, Cytokines, and Oxidative Stress. Front Microbiol. 2021;12:656372. 10.3389/fmicb.2021.656372

65. Krishnareddy S. The Microbiome in Celiac Disease. Gastroenterol Clin. Elsevier; 2019;48:115–26. 10.1016/j.gtc.2018.09.008

66. Valitutti F, Cucchiara S, Fasano A. Celiac Disease and the Microbiome. Nutrients. Multidisciplinary Digital Publishing Institute; 2019;11:2403. 10.3390/nu11102403

67. Prins FM, Collij V, Groot HE, Björk JR, Swarte JC, Andreu-Sánchez S, et al. The gut microbiome across the cardiovascular risk spectrum. Eur J Prev Cardiol. 2024;31:935–44. 10.1093/eurjpc/zwad377

68. Vandeputte D, De Commer L, Tito RY, Kathagen G, Sabino J, Vermeire S, et al. Temporal variability in quantitative human gut microbiome profiles and implications for clinical research. Nat Commun. Nature Publishing Group; 2021;12:6740. 10.1038/s41467-021-27098-7

69. Gajer P, Brotman RM, Bai G, Sakamoto J, Schütte UME, Zhong X, et al. Temporal Dynamics of the Human Vaginal Microbiota. Sci Transl Med. American Association for the Advancement of Science; 2012;4:132ra52–132ra52. 10.1126/scitranslmed.3003605

70. Gerber GK. The dynamic microbiome. FEBS Lett. 2014;588:4131–9. 10.1016/j.febslet.2014.02.037

